# Measuring ontogenetic shifts in central-place foraging insects: a case study with honey bees

**DOI:** 10.1101/2020.03.31.017582

**Authors:** Fabrice Requier, Mickaël Henry, Axel Decourtye, François Brun, Pierrick Aupinel, François Rebaudo, Vincent Bretagnolle

**Author notes:** Corresponding author: Fabrice Requier.

## Abstract

1. Measuring time-activity budgets over the complete individual lifespan is now possible for many animals with the recent advances of life-long individual monitoring devices. Although analyses of changes in the patterns of time-activity budgets have revealed ontogenetic shifts in birds or mammals, no such technique has been applied to date on insects.
2. We tested an automated breakpoint-based procedure to detect, assess and quantify shifts in the temporal pattern of the flight activities in honey bees. We assumed that the learning and foraging stages of honey bees will differ in several respects, to detect the age at onset of foraging (AOF).
3. Using an extensive dataset covering the life-long monitoring of 2,100 individuals, we compared the AOF outputs with the more conventional approaches based on arbitrary thresholds. We further evaluated the robustness of the different methods comparing the foraging time-activity budget allocations between the presumed foragers and confirmed foragers.
4. We revealed a clear-cut learning-foraging ontogenetic shift that differs in duration, frequency, and time of occurrence of flights. Although AOF appeared to be highly plastic among bees, the breakpoint-based procedure seems better able to detect it than arbitrary threshold-based methods that are unable to deal with inter-individual variation.
5. We developed the *aof* R-package including a broad range of examples with both simulated and empirical dataset to illustrate the simplicity of use of the procedure. This simple procedure is generic enough to be derived from any individual life-long monitoring devices recording the time-activity budgets of honey bees, and could propose new ecological applications of bio-logging to detect ontogenetic shifts in the behaviour of central-place foraging insects.

## Introduction

Recent advances in automated life-long individual monitoring devices revolutionized data collection and accessible information on free-ranging animals. The automated record of time-activity budgets has especially provided unprecedented perspectives on central-place foragers (Ropert-Coudert & Wilson, 2005; Rishworth, Tremblay, Green, & Pistorius, 2014). Central-place foraging –defined by successive trips between a central-place and foraging patches (Orians & Pearson, 1979)– is a fairly common foraging strategy used by animals that are constrained at some point of their life in a central place. Many species adopt this foraging strategy, such as birds flying from a nest or a colony, mammals walking from a burrow (e.g. voles), flying from a cavity or a colony (e.g. bats), or diving from the coast (e.g. sea lions), and insects foraging from a nest or a colony (e.g. ants, wasps, bees). While it is well established that recording the patterns of time-activity budgets in central-place foraging can provide information on foraging traits such as habitat use and diet (e.g. Lamb, Satgé, & Jodice, 2017; Briscoe et al., 2018), fewer studies have revealed their potential for analysing keystone ontogenetic traits (e.g. Lambin & Yoccoz, 2001; Gopukumar, 2003; Benton, St Clair, & Plaistow, 2008; Fay, Barbraud, Delord, & Weimerskirch, 2016).

The detection of changes in patterns of time-activity budgets may reveal ontogenetic shifts, such as parental age (Fay et al., 2016), maternal age (Benton et al., 2008) and childbirth age (i.e. the age in between pregnancy and lactation stages in mammals, Lambin & Yoccoz, 2001; Gopukumar, 2003). Ontogenetic shifts are critical within an evolutionary context; early ontogenetic transition may affect growth, mortality, and therefore individual fitness (Post, 2003). However, bio-logging applications are still restricted to birds (particularly seabirds, e.g. Rishworth et al., 2014; Fay et al., 2016; Lamb et al., 2017), fishes (Gurarie et al., 2016) and mammals (Lambin & Yoccoz, 2001; Gopukumar, 2003; Gurarie, Andrews, & Laidre, 2009; Briscoe et al., 2018). Very few studies were conducted on insects (but see Degen et al., 2016 for bees), despite the recent availability of adapted miniaturized devices (Kissling, Pattemore, & Hagen, 2014; Nunes-Silva et al., 2019). Here we examine whether the use of life-long individual monitoring devices can reveal the ontogenetic shifts in the behaviour of the free-ranging western honey bees *(Apis mellifera* L.), as a case study of central-place foraging insects.

Honey bees are social insects living in a colony nest. Foraging tasks are carried out by a very reduced subset (around 7 %) of the adult worker population by successive round trips between the nest and flower patches (Winston, 1994). The honey bee presents the advantage of living in a single nest composed of a very large number of individuals (usually several tens of thousands of individuals, Winston, 1994), and a rather short lifespan (usually several tens of days, Winston, 1994), which facilitates life-long monitoring of many individuals simultaneously. To date, three main techniques have been used to monitor time-activity budgets in honey bees (and other small-sized insects), the harmonic radar (Capaldi et al., 2000), the Radio-Frequency IDdentification device (RFID; Streit, Bock, Pirk, & Tautz, 2003) and the image-based optical counter (Chen, Yang, Jiang, & Lin, 2012). With such devices, the time-activity budgets of flights (duration and number of trips per day) are recorded at the individual level.

One main critical aspect of life-history in honey bees are the ontogenetic shifts at the adult stage (Visscher & Dukas, 1997; Becerra-Guzman, Guzman-Novoa, Correa-Benitez, & Zozaya-Rubio, 2005; Rueppell, Bachelier, Fondrk, & Page, 2007; Perry, Søvik, Myerscough, & Barron, 2015; Klein et al., 2019). The “Age at Onset of Foraging” (AOF) –defined as the age at the onset of foraging activity *per se* when adult bees have completed the learning phase occurring during orientation flights (Capaldi, & Dyer, 1999; Capaldi et al., 2000; Degen et al., 2016)– is arguably a critical fitness trait of honey bees at both individual and colony levels (Perry et al., 2015). An earlier ontogenetic transition to foraging reduces the foraging performance of honey bees at individual level, i.e. the number of days as a forager (Visscher & Dukas, 1997; Becerra-Guzman et al., 2005; Rueppell et al., 2007; Perry et al., 2015) and can impacts the fitness of the whole colony (Becher et al., 2014; Perry et al., 2015). However, although critical, AOF is challenging to detect in the field. Previous studies of flight movement of free-ranging honey bees suggested possible detections of AOF using patterns of time-activity budgets. Indeed, learning flights are concentrated near the hive, within a few hundred meters (median of 189m in Capaldi et al., 2000) while foraging flights take place over much greater distances (about 1.7 km on average; Beekman & Ratnieks, 2000; Steffan-Dewenter & Kuhn, 2003), presumably transferring into flight time duration. Hence the duration and the number of trips, as proxies of time-activity budgets, should differ between learning and foraging, and AOF was thus commonly defined using arbitrary thresholds. For instance, ontogenetic shift was defined as the first day that a bee performs 5 trips or performs a trip longer than 30 minutes (Calderone & Page, 1988; Dukas, 2008, Tenczar, Lutz, Rao, Goldenfeld, & Robinson, 2014; He, Tian, Wu, & Zeng, 2015; Perry et al., 2015; Klein et al., 2019). However, the use of arbitrary thresholds may fail to capture the true timing of ontogenetic shifts, given the high inter-individual variation in honey bee life-history traits (Huang & Robinson, 1996; Amdam et al., 2009; Perry et al., 2015), and may differ with the genetic composition of the colony (Calderone & Page, 1988; Becerra-Guzman et al., 2005), climatic environmental factors (Henry et al., 2014; He et al., 2015), landscape structure and composition (Henry et al., 2014), and pesticide exposure (Decourtye et al., 2011; Henry et al., 2012; Prado et al., 2019). Moreover, broad behavioural plasticity among honey bees from the same cohort (i.e. bees with same birth date; Tenczar et al., 2014) suggests the presence of inter-individual variation even in the same genetic and environmental contexts.

Assuming that time-activity budgets of learning and foraging flights of honey bees would differ either in duration, frequency, or time of occurrence in the day (Capaldi et al., 2000), we developed a simple statistical procedure, (*i*) working at the individual level thus accounting for inter-individual variation, to (*ii*) detect, *(iii)* assess and *(iv)* quantify shifts in the temporal pattern of time-activity budgets recorded by individual life-long monitoring. We based the statistical procedure on the behavioural change point analysis approach (BCPA; Gurarie, Andrews, & Laidre, 2009), a well appreciated technique of likelihood comparisons to statistically determine behavioural changes. BCPA is particularly acknowledged for the ‘window sweep’ function determining where along an animal’s trajectory changes in the behavioural state occur based on changes in the underlying movement patterns (Gurarie et al., 2009; 2016; Kranstauber et al., 2012; Zhang et al., 2015). However, this complex function was developed for time-stamped movement data, i.e. in which there are X, Y and T coordinates representing spatial locations and time of observation, rather than for simple time series data (Gurarie 2013). Here, we considered the simplest ‘GetBestBreak’ function –a BCPA function developed for detecting a single behavioural change in irregular univariate time series (Gurarie 2013)– for its application to measure ontogenetic shifts in honey bees. Then we tested the statistical procedure on an extensive RFID dataset covering the life-long monitoring of 2,100 individual honey bees. We further compared our AOF outputs with the more conventional approaches based on arbitrary thresholds, evaluated the robustness of the procedure with different types of datasets, and compared the foraging time-activity budget allocations between the predicted foragers and observed foragers. We also explored the relationship between AOF ontogenetic plasticity and foraging performance in honey bees, and we tracked the potential risk of early ontogenetic transition over time. Finally, we provide guidelines (the *aof* R-package) on how using the procedure to detect life-history transitions of free-ranging insects from time series data, as a generic approach towards measuring ontogenetic shifts in central-place foraging insects.

## Material and methods

### STUDY SITE & HONEY BEE COLONIES

The study was performed in the Long Term Social-Ecological Research “Zone Atelier Plaine & Val de Sèvre” (LTSER ZA-PVS) in central western France, 46°23’N-0°41’W (see Bretagnolle et al., 2018 for a general presentation). This study area has hosted the ECOBEE platform since 2008, a field-realistic observatory of managed honey bee colonies investigating population ecology at the landscape scale (Odoux et al., 2014). In 2011, three honey bee colonies from a livestock managed apiary of *A. mellifera mellifera × caucasica* strain at INRA Le Magneraud (46°09’N, 0°41’W, about 30 km west of the study area) were placed in the study area (**Supporting Information, Figure S1**). Queens were one year old and their health status was checked for any visible disease symptoms. The hive model was the common Dadant-Blatt model (with 10 brood frames) in pine wood-waxed microcrystalline.

### LIFE-LONG INDIVIDUAL MONITORING DEVICE

Individual honey bees were monitored using RFID tags (mic3^®^ – TAG 64 bit RO, iID2000, 13.56 MHz system, 1.0 mm *×* 1.6 mm *×* 0.5 mm; Microsensys GmbH, Erfurt, Germany) and ad-hoc readers (iID2000, 2k6 HEAD; Microsensys GmbH, Erfurt, Germany) placed at the entrance of the hives (**Figure 1a**). Readers were powered by a turnover of two slow-discharge rate batteries (12V, 92Ah, C20, Banner®). The readers function as data loggers, recording the identity, date, and time (to the nearest 1 second) of each tag detection, also termed *hit* (tag-reader detection range = 3 mm; see also Streit et al., 2003). The readers’ internal clock synchronisation and daily data download were automatically processed by a permanently connected laptop using custom-made software (BeeReader, Tag Tracing Solution, Valence, France). Hits from the different readers were then collected and coded into a unique ASCII text file.

**Figure 1.**
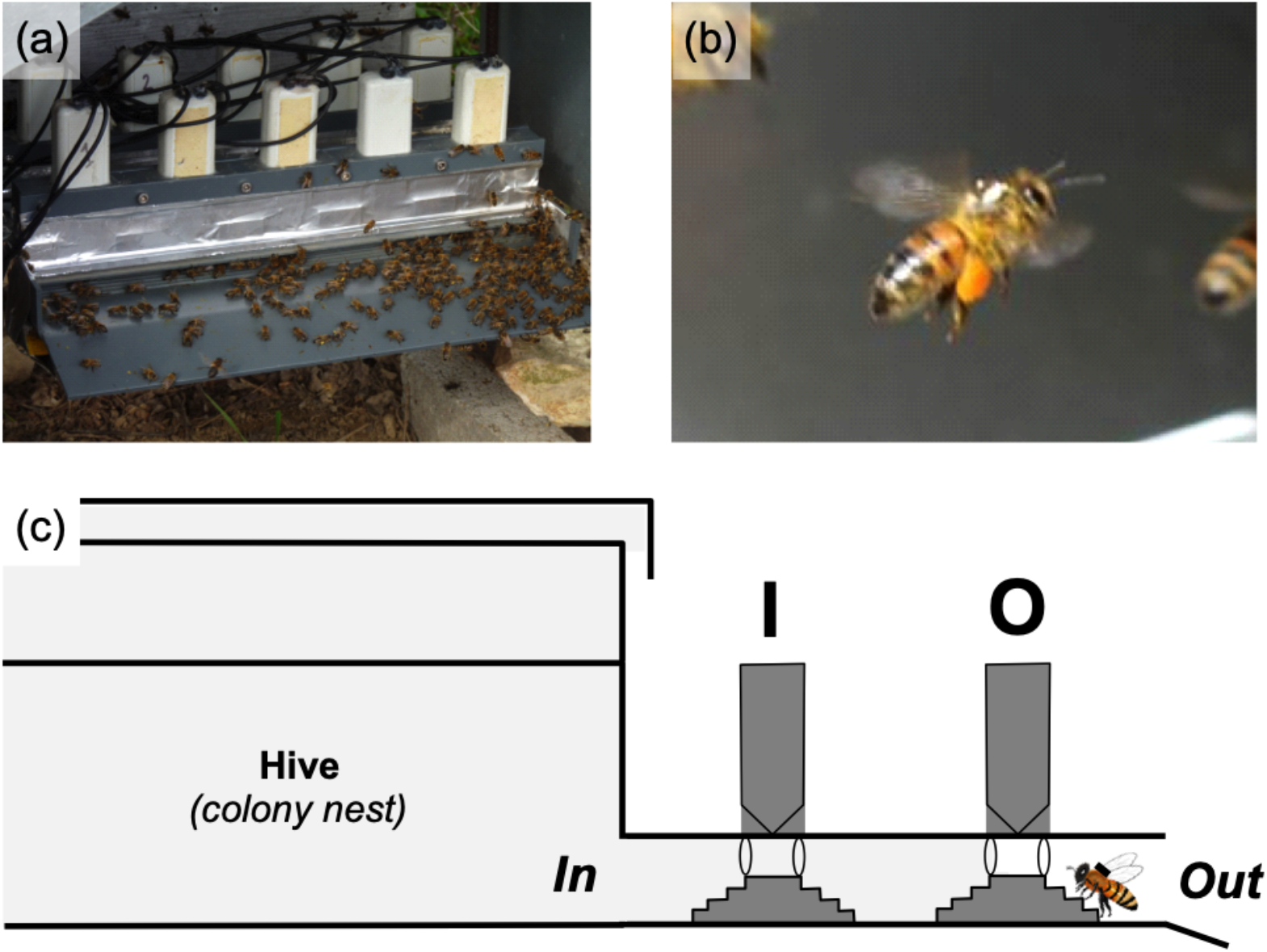
Honey bee RFID monitoring device adapted to common Dadant beehives. (a) The hive entrances were equipped with two rows of five RFID-readers in order to discriminate in-and-out activities of honey bees. (b) A pollen-forager honey bee equipped with RFID-tag. (c) The conceptual schema of RFID adaptation for detecting in-and-out activities of honey bees throughout a free-walking room (i.e. the room between the two rows of RFID-readers I and O). Bees were strictly forced to pass under a I reader and a O reader for an exit activity, and vice-versa, allowing the complete sequencing of all the in-and-out activities. To limit the risk of in-and-out activity congestion due to readers, bees were able to stay between the two rows of reader, nevertheless, they were systematically recorded when leaving this transition zone.

RFID tags weighed approx. 3 mg, i.e. 3 % of an adult honey bee’s body mass. This weight is considered low enough to not interfere with the individual life and tasks (Streit et al., 2003), inasmuch as honey bees typically carry pollen and nectar loads representing 20% and 35% of body mass, respectively, maximally reaching 80% of body mass (Winston, 1994, **Figure 1b**). Tags were fitted on the bee’s thorax using biocompatible dental cement (TempoSIL®2, Coltène/Whaledent s.a.r.l., Le Mans, France). The full description of applying the RFID on bees may be found in Decourtye et al. (2011).

### MEASURING TIME-ACTIVITY BUDGETS

During the entire experiment, a total of 2,100 newly emerged sister workers (one day-old) were collected from a single honey bee colony at the INRAE Entomology experimental unit livestock apiary, and processed as part of six successive cohorts evenly staggered from April to September of 2011 (**Table S1**). Each honey bee was equipped with a unique RFID tag, and released in cohorts of 75-150 individuals into the three experimental colonies (**Table S1**).

The time-activity budgets were recorded using two adjacent rows of five contiguous RFID readers placed at the entrance of the hives (i.e. the common 10-frame Dadant hive, **Figure 1a**). Bees were strictly forced to pass under the readers for all their in-and-out activities (i.e. no other entrance to the nest was available). The inner reader row (I) and outer row (O) are identified along with each hit, making it possible to distinguish between incoming and outgoing flights (O-I *vs.* I-O hit successions, respectively, **Figure 1c**). The sequences *(seq)* of incoming and outgoing flights delineate the time periods spent in-nest or outside of the hive, providing the opportunity to investigate time-activity budgets. We relied on the “Track a forager” method (Van Geystelen, Benaets, de Graaf, Larmuseau, & Wenseleers, 2016) to infer time-activity budgets from the raw RFID dataset. We modified the analytical process (**Section S1**) to adapt the method to the common 10-frame Dadant hive by establishing two rows of five RFID readers over the entire width of the hive entrance, and free-walking room between the two rows of readers. This standard process provides essential time-activity budget variables from RFID tracking, namely the number of trips per day *(Trip number),* the mean duration of trips per day *(Trip duration,* in seconds), and the average time of day of the trip *(Trip time,* in hours).

### DECISION RULES TO DETERMINE ONTOGENETIC SHIFTS

Our aim was to find an automated method to assess the Age at Onset of Foraging (AOF), a well-established ontogenetic shift in honey bees, which marks a change in flight time-activity budgets between learning flights and foraging flights. For that purpose, we used our three candidate time-activity budget variables: *Trip number, Trip duration,* and *Trip time.* The latter parameter was chosen because orientation/learning flights are performed later in the day compared to foraging flights that are more evenly distributed during the daytime (Tanczar et al., 2014). The time of day of a given trip was represented by the time at the beginning of the trip. We then ran the BCPA *GetBestBreak* R-function (Gurarie 2013) to detect a single behavioural change in univariate time series. We tested the parsimony of the breakpoint prediction using the ‘GetModels’ function. Following Gurarie (2013), we considered the predicted breakpoint as parsimonious whenever Δ *Bayesian information criterion* (BIC) > 2 between the simple model (without breakpoint) and the complete model (**Figure 2a**). Successful detections were pooled in the “detected” group, and unsuccessful ones in the “undetected” group. Individuals with no trip recorded (i.e. only one hit) were discarded from the analysis as well as the individuals with less than five days with recorded trips because breakpoints could not be established; both were pooled in the “uncomputed” group. All statistical analyses were performed using the R Project for Statistical Computing version 3.6.2 (R Development Core Team, 2020). To facilitate the use of this procedure by other researchers, we provide an *aof* R-package with examples using both simulated and empirical dataset. Documentation and source code are freely available on CRAN (https://cran.r-project.org/package=aof) and GitHub (https://github.com/frareb/aof). We then averaged the values of breakpoint age obtained from two candidate time budget variables, *Trip duration* and *Trip time,* to provide the AOF estimates. *Trip number* was discarded because of the non-parsimonious estimation of its AOF value *(Trip time* was considered due to marginally significant effect, see *results).* We *a posteriori* assessed the overall difference in time budget allocations between the two life stages, learning and foraging, using Generalized Linear Mixed Models (GLMMs) with the *lme* function in the *nlme* R-package. We specified a Gaussian error structure and the origin of each honey bee (crossed cohort × site grouping) as a context random effect.

**Figure 2.**
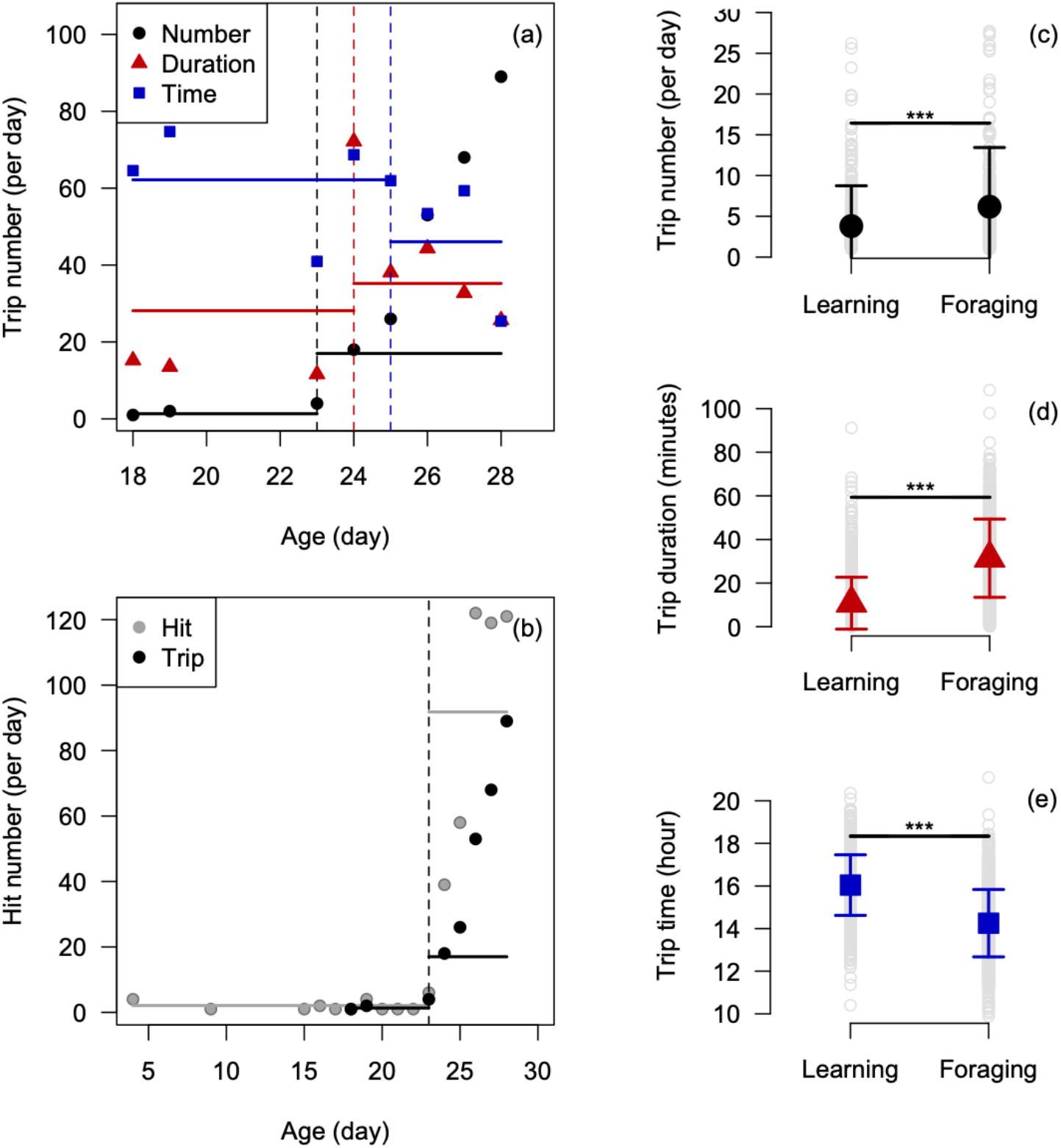
Breakpoint-based method to estimate the Age at the Onset of Foraging (AOF) in honey bees from time-activity budgets. (a) The daily activities was computed at the individual level (here the bee *#A00103C00040A5F4)* from each time budget variable on trips (i.e. number, duration and time of trips). The best candidate breakpoint age gives the AOF estimate (vertical lines). (b) The described breakpoint-based method can also be applied directly from the output of the formatting phase, to search among the daily cumulative hit numbers (illustrated with the same bee *#A00103C00040A5F4).* The proposed breakpoint-based method estimates AOF as a change in flight time-activity budget between learning flight and foraging flight periods in honey bees, with differences in (c) the number of trips per day; n = 492 successful AOF detected (i.e. “detected” bees), (d) the mean duration of trips per day; n = 835, and (e) the average time of day of a trip in hours; n = 835. Dots show the mean ± SE of the time-activity budget variables.

### ASSESSMENT OF ROBUSTNESS OF ONTOGENETIC SHIFT DETECTION

The resulting breakpoint-based estimates of the AOF ontogenetic shift were first compared with values computed with raw RFID data, the latter allowing the use of missing detections and subsequent disturbances in time-activity budget estimates. Indeed, individual monitoring devices commonly harbour missing detections, particularly with small species (i.e. ants, bees, wasps; see Streit et al., 2003; Tanczar et al., 2014; Van Geystelen et al., 2016), disturbing the estimation of time-activity budget variables resulting from the sequencing process. Consequently, whenever the detection bias cannot be firmly established or is expected to be low, it might be safer to use an alternative AOF measurement approach that does not rely on the assumption of unaltered trip sequences. Herein, we applied a breakpoint detection procedure based on hits rather than trips. In other words, we directly used the output of the raw ‘hits’ from the RFID readers, thereby skipping the sequencing phase (**Section S1**) to search among the daily cumulative hit numbers *(Hit number)* and the average time of day of the hits (*Hit time,* in hours).

We also compared the resulting breakpoint-based estimates of AOF with values obtained from the more traditionally used, but arbitrary, threshold approaches. We considered two options: AOF defined either (*i*) as the age when the bee performed for the first time more than 5 trips per day (e.g. Tenczar et al., 2014) or (*ii*) as the age when the bee performed for the first time flight durations longer than 30 minutes (e.g. Perry et al., 2015).

Finally we compared the foraging time-activity budget allocations (i.e. number, duration and time of trips) between confirmed (observed) foragers and presumed (predicted) foragers, to evaluate the robustness of the method. Observed foragers were deduced from the presence of pollen fixed on their hind legs when at the entrance of the hive. This visual point mark allowed us to collect data on 67 foragers on the 21^st^ of June 2011. We equipped these confirmed foragers with RFID and reintroduced them into their original colony (the hive A, **Table S1**). We compared their foraging time-activity budget allocations with 75 presumed foragers, i.e. emerged bees equipped with RFID and introduced into the same colony 20 days before the reintroduction of the confirmed foragers (**Table S1**), the average time required for emerged bees to perform their first foraging activity (Winston, 1994). We then applied the breakpoint-based procedure to detect the AOF of presumed foragers and their foraging time-activity budget allocation. We used a *t*-test to statistically assess the mean difference in foraging time-activity budget allocations between presumed foragers and confirmed foragers. We finally computed the Cohen’s *d* value (*tes* function in the *compute.es* R-package) to compare the magnitude of difference in effect sizes between the candidate time budget variables, ranging from 0, i.e. no difference, to 1, i.e. a marked difference (Furukawa & Leucht, 2011).

### INTEGRATING ONTOGENETIC SHIFTS IN HONEY BEE LIFE-HISTORY

Life-history may be defined as the timing of occurrence and duration of different life stages. For example, honey bees have an in-nest task stage before the Age at First Exit (AFE), a learning flight stage before AOF, and a foraging stage until death (LSP for lifespan). Most honey bees die outside of the hive, because foraging bees are exposed to increased mortality risks, including predation, disorientation and weather hazards (Visscher & Dukas, 1997). In the opposite case of death inside the hive, workers carry the dead counterparts out of the hive as soon as possible to preserve the sanitary state of the colony (Winston, 1994). AFE was defined as the age (days) at the first trip record, i.e. the first OO chain with > 4s interval. AFE provides information on the duration of time that an individual honey bee was assigned to innest tasks. LSP is obtained by the age (days) at the last hit.

We explored the relationship between the AOF ontogenetic plasticity and foraging performance by computing the relationships between (1) AOF and foraging stage (i.e. the number of days of foraging as the difference between LSP and AOF) and (2) AOF and the number of foraging trips (log_10_ transformed), using GLMMs with Gaussian error structure and the origin of each honey bee (crossed cohort × site grouping) as a contextual random effect. We computed both linear and quadratic models and compared them using the using the *Akaike Information Criterion* (AIC) value. When a non-linear quadratic relationship was suspected (ΔAIC > 2) with an intermediate AOF value maximizing the foraging performance, the optimal AOF value was estimated as the peak of the quadratic curve. Finally we analysed the seasonality of AOF to explore potential strategies of *precocity* vs. *lateness* in bees, by investigating AOF that occur before *vs.* after the optimal AOF value, from April to September.

### SENSITIVITY ANALYSIS OF ONTOGENETIC SHIFT DETECTION

Although the procedure was developed to detect the Age at Onset of Foraging (AOF) in honey bees, this method could be of interest for other species. Thus, we carried out a sensitivity analysis by simulating a broad range of virtual data to assess the relevance and limit of our procedure. We first simulated two sets of data: (*i*) a “no change simulated” scenario to assess the sensitivity of the procedure to detect behavioural changes when there are none, and (*ii*) a “change simulated” scenario to assess the robustness of the procedure to detect existing behavioural change (i.e. a single breakpoint in time series). Both scenarios included time series from 1 to 50 days with a breakpoint at day 25 in case of the “change simulated” scenario (the “no change simulated” scenario had a single *μ* value *[μ* = 50] over the time series). The breakpoint was provided by alteration of the *μ* value between the two stages, with *μ* = 25 in stage 1 (before day 25) and *μ* = 50 in stage 2 (after day 25) (**Section S2**).

Assuming that the performance of the procedure to correctly detect (or not) behavioural changes could be affected by the number of data points in time series and the variance in data, we additionally modified these two parameters in both scenarios. For that, we created a factorial design for which we sequentially increased the number of data points (*n*) from 5 to 45 over 40 categories, and we sequentially increased the variance (*v*) from *σ* = 0.01 to *σ* = 3 over 40 categories (**Section S2, Figure S3**). We then transformed v as percentage of variance simulated from 0 % to 100 %, respectively. Thus, we simulated at set of 1600 data (40 *n* categories × 40 *v* categories) for the “no change simulated” scenario and 1600 data for the “change simulated” scenario.

To assess the sensitivity of the procedure, we analysed whether the detection rate of behavioural changes varied with the number of data points (*n*) and the data variance (*v*). For that, we fit a generalized linear model (GLM) with binomial error structure (*glm* function in the *base* R-package). The model included both variables (*n* and *v)* as quantitative predictors, and their interaction. We used the same modelling approach for analysing the robustness of the procedure to detect existing behavioural changes.

## Results

### ONTOGENETIC SHIFTS IN HONEY BEES

From the 2,100 tagged honey bees, 11.1 % (233 bees, on average 15.3 ± 8.1 per cohort) did not provide any data and were discarded from the analysis. For the rest, we collected a total of 402,471 *hits,* yielding 256,462 *seq* and 21,104 *trips* from 1,867 bees, over a non-stop monitoring record of 193 days. Time-activity budgets were used to estimate the age at onset of foraging, AOF. Excluding 31.1 % of bees which provided only one *hit,* and 6.4 % providing less than 5 trips, which were both pooled in the “uncomputed” group, a breakpoint could be identified in 42.2 to 88.5 % of the 1,167 remaining bees (depending of the time budget variable considered, **Supporting Information Section S3**), either from daily trip numbers, trip durations or average time of trips in the day (**Figure 2a**).

GLMMs performed *a posteriori* (i.e. once the breakpoint was detected) indicated that the AOF significantly differentiated two life-history stages, learning and foraging, from their time-activity budget allocations (**Figures 2c-e**). Compared to the learning stage, foragers performed more trips per day (GLMM with cohort as random effect, *F*_1,491_ = 54.38, *p* < 0.001, *n* = 492, **Figure 2c**), longer trip durations (*F*_1,834_ = 1,023.79, *p* < 0.001, n = 835, **Figure 2d**) and at a slightly but significantly earlier average time during the day (*F*_1,834_ = 618.51, *p* < 0.001, n = 835, **Figure 2e**).

### ASSESSMENT OF ROBUSTNESS OF ONTOGENETIC SHIFT DETECTION

Overall, estimates of the AOF ontogenetic shift differed by only 0.1 ± 1.1 to 2.5 ± 1.1 days (mean ± SE) according to the different methods and time budget variables, whether they were the breakpoint-based methods on trips, the breakpoint-based methods on hits, or the arbitrary threshold-based methods (**Section S3**). These uncertainties are rather low considering they represent only 0.3 – 8.1 % of the average total lifespan (31.0 days). For instance, the breakpoint-based method relying on hit *vs.* trip numbers differed by 2.4 ± 1.2 days (**Figure 2b, Section S3**). However, they differed far more with regard to the successful detection of AOF (from 492 to 1,033 bees) and the similarities in the identity of detected bees (the proportion of shared detected bees between pairwise candidate time budget variables ranged from 42.4 ± 10.0 % to 94.8 ± 4.0 %, **Section S3**). Overall, the breakpoint-based methods using hit number and hit time provided more detections while the breakpoint-based methods using trip duration and trip time provide very similar estimates (both in term of AOF date and identity similarity of bees with detection; **Section S3**). Conversely, the breakpoint-based method using trip number performed the least well, with low detection rates and similarity, i.e. sharing a low proportion of detected bees with other time budget variables (**Section S3**).

Comparing foraging time-activity budgets between confirmed (n = 67) and presumed foragers (n = 39 bees with AOF detected) confirmed the decreased performance of arbitrary threshold-based methods (**Figure 3**). While no significant differences arose for breakpoint-based methods (i.e. *Hit number, Hit time* and *Trip duration),* arbitrary threshold-based methods showed significant differences (*t* = 4.78, *p* < 0.001 for the threshold-based method using 5 trips per day, and *t* = −2.42, *p* = 0.019 for the threshold-based method using flight durations longer than 30 minutes; **Table S2**). The breakpoint-based method using *Trip number* also showed a significant difference (*t* = 3.30, *p* = 0.003) as well as *Trip time* to a lesser extent (*t* = – 2.33, *p* = 0.023; **Table S2**), suggesting that *Trip number* and *Trip time* should be discarded as methods to estimate AOF in bees (**Figure 3**).

**Figure 3.**
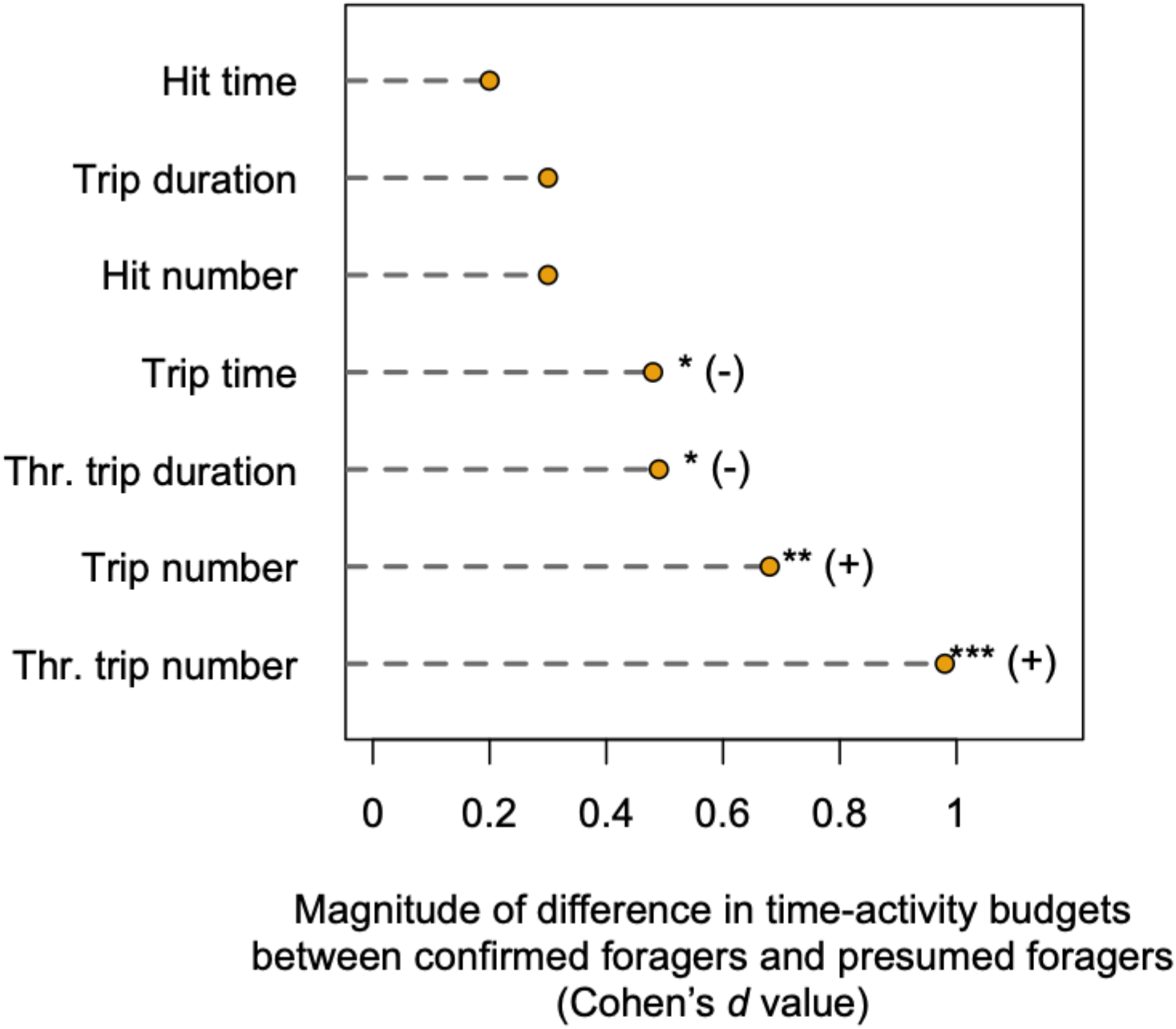
Comparison in the foraging time-activity budget allocations (i.e. number, duration and time of trips) between confirmed foragers (n = 67) and presumed foragers (n = 39 bees with AOF detected). *Thr. trip number* means the threshold-based method using 5 trips per day, and *Thr. trip duration* means the threshold-based method using flight durations longer than 30 minutes. The other time budget variables involve the breakpoint-based method on hit or trip. The mean difference in foraging time-activity budget allocations was expressed with the Cohen’s d value to compare the magnitude of difference in effect between the candidate time budget variables, ranging from 0, i.e. no difference, to 1, i.e. large difference. *P*-values (i.e. statistics of the *t*-test) and the directions of the differences were also expressed. *** = *p* < 0.001; ** = *p* < 0.01; * = *p* < 0.05; (-) = underestimation of the method; (+) overestimation of the method.

### INTEGRATING ONTOGENETIC SHIFTS INTO HONEY BEE LIFE-HISTORY

Using a single AOF value per individual, computed as the averaged AOF among the two breakpoint-based methods *(Trip duration* and *Trip time;* the latter was considered due to marginally significant effect, see above), the average life-history of honey bee workers (**Figure S2**) was characterized by a mean lifespan of 31.0 ± 10.2 days, encompassing an initial in-nest stage lasting 12.2 ± 6.9 days (as given by AFE), a learning flight stage lasting 9.4 ± 7.4 days, and a final foraging stage lasting 8.7 ± 4.1 days with an AOF occurring at 22.4 ± 10.6 days old (n = 865, corresponding of 74.1 % of successful AOF detected). During the foraging stage, honey bees performed an average of 6.2 ± 7.3 trips per day (**Figure 2c**), with flight durations of about 31.4 ± 17.9 minutes (**Figure 2d**). However, these values presented a high inter-individual variability (e.g. 1 to 28 foraging trips per day, or 0.1 to 119.0 min for the foraging flight duration per trip). In addition, all bees did not perform all stages (i.e., in-nest tasks + learning + foraging): 31.1 % of the monitored bees did not perform outside stages (corresponding to the “no computed” group, n = 653) though an unknown proportion may correspond to individuals dying in nest.

There was a non-linear relationship between AOF and the number of foraging days (GLMM with cohort as random effect, *F*_3,862_ = 24.771, *p* < 0.001; **Figure 4a**). While the time lag between AOF and LSP, the so-called foraging performance, was 8.7 ± 4.1 days (**Figure S2**), AOF affected this duration quadratically rather than linearly (ΔAIC = 22.4 between quadratic and linear). The duration of the foraging stage increased gradually from an AOF of 6 days to an optimal AOF value of 41 days (the peak of the quadratic curve), i.e. the AOF age that maximizes the duration of the foraging stage. The duration of the foraging stage gradually decreased from this optimal AOF value to 60 days (**Figure 4a**). The relationship between AOF and the number of foraging trips also showed a non-linear (although less marked), quadratic (ΔAIC = 2.46 between quadratic and linear), pattern (*F*_3,862_ = 4.468, *p* < 0.001; **Figure 4b**). Moreover, AOF showed a high seasonal variation from April to September (**Figure 4c**), starting around 18 days from April to June in all colonies, increasing to approximately 35 days in July, then decreasing again to 20 days in August and September (**Figure 4c**).

**Figure 4.**
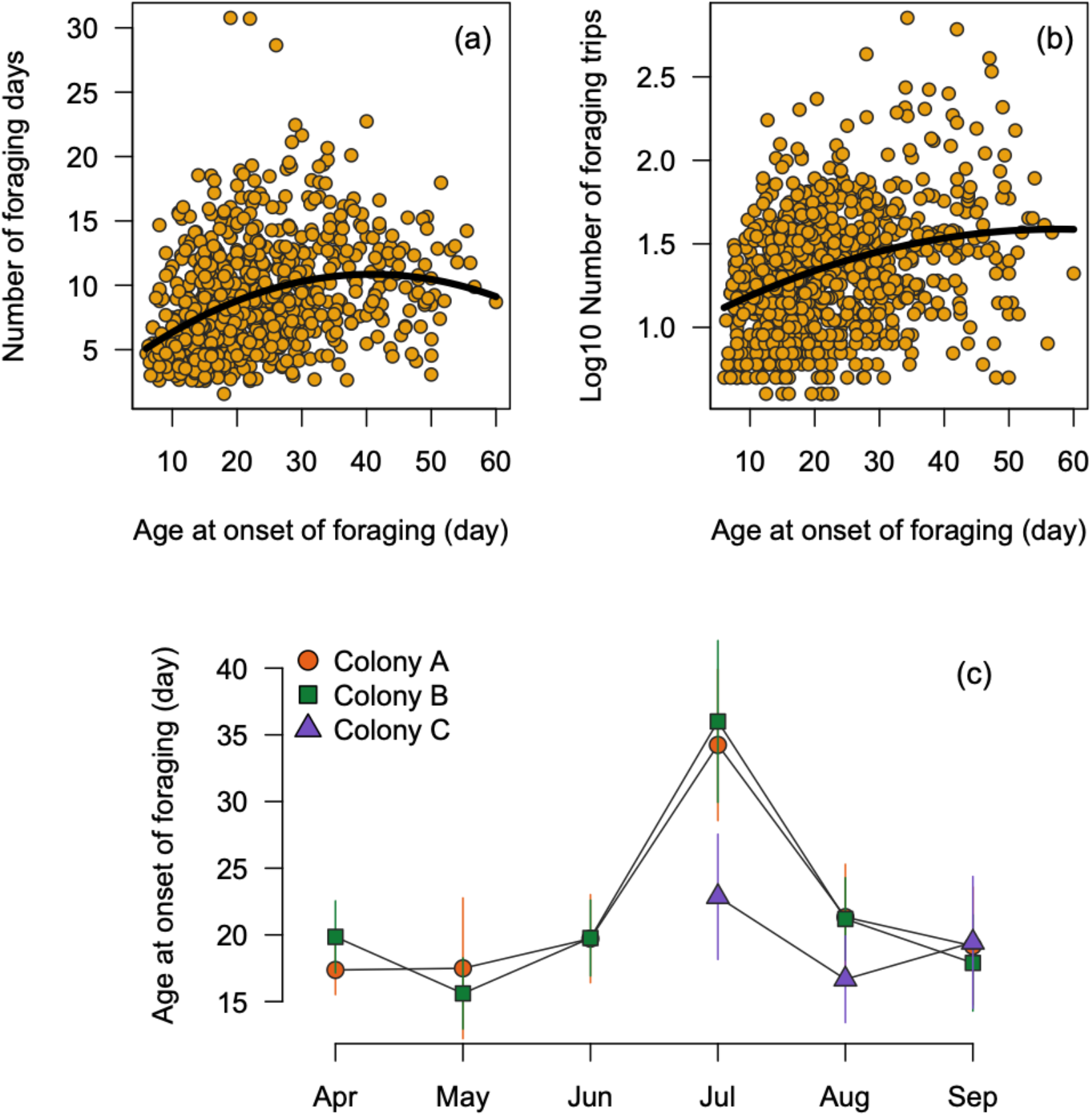
Relationship between ontogenetic plasticity and foraging performance in honey bees. (a) The non-linear relationship between the age at onset of foraging (AOF) and the foraging stage highlighted an optimal AOF value maximizing the foraging performance in bees at AOF = 41 days. (b) The relationship between AOF and the number of foraging trips also showed a non-linear, quadratic, pattern. (c) The temporal dynamic of the AOF value from April to September indicated a high seasonal variation with a general tendency of precocity in bees (i.e. AOF below 41 days). Dots show the mean ± 1/2SE of the AOF estimate.

### SENSITIVITY ANALYSIS OF ONTOGENETIC SHIFT DETECTION

Although the automated breakpoint-based procedure was initially developed to detect AOF in honey bees, it may be of interest for measuring other behavioural changes in other species (e.g. ants, wasps, **Figure 5a**). The false detection rate of behavioural changes was very low (0.06 for null data variance) in the *No change simulated* scenario (intercept value, *Z* = −7.519, *p* < 0.001, **Figure 5b**) and very high (0.83 for 10 data points) in the *Change simulated* scenario (intercept value, *Z* = 3.940, *p* < 0.001, **Figure 5c, Table S3**). The false detection rate substantially increased with variance in the data (*Z* = 6.980, *p* < 0.001), an effect partially buffered by the number of data points (**Figure 5b, Table S3**). Otherwise, the detection rate of true behavioural changes *(Change simulated* scenario) increased with the number of data points *(Z* = 2.587, *p* = 0.010), an effect mitigated by the data variance (**Figure 5c, Table S3**).

**Figure 5.**
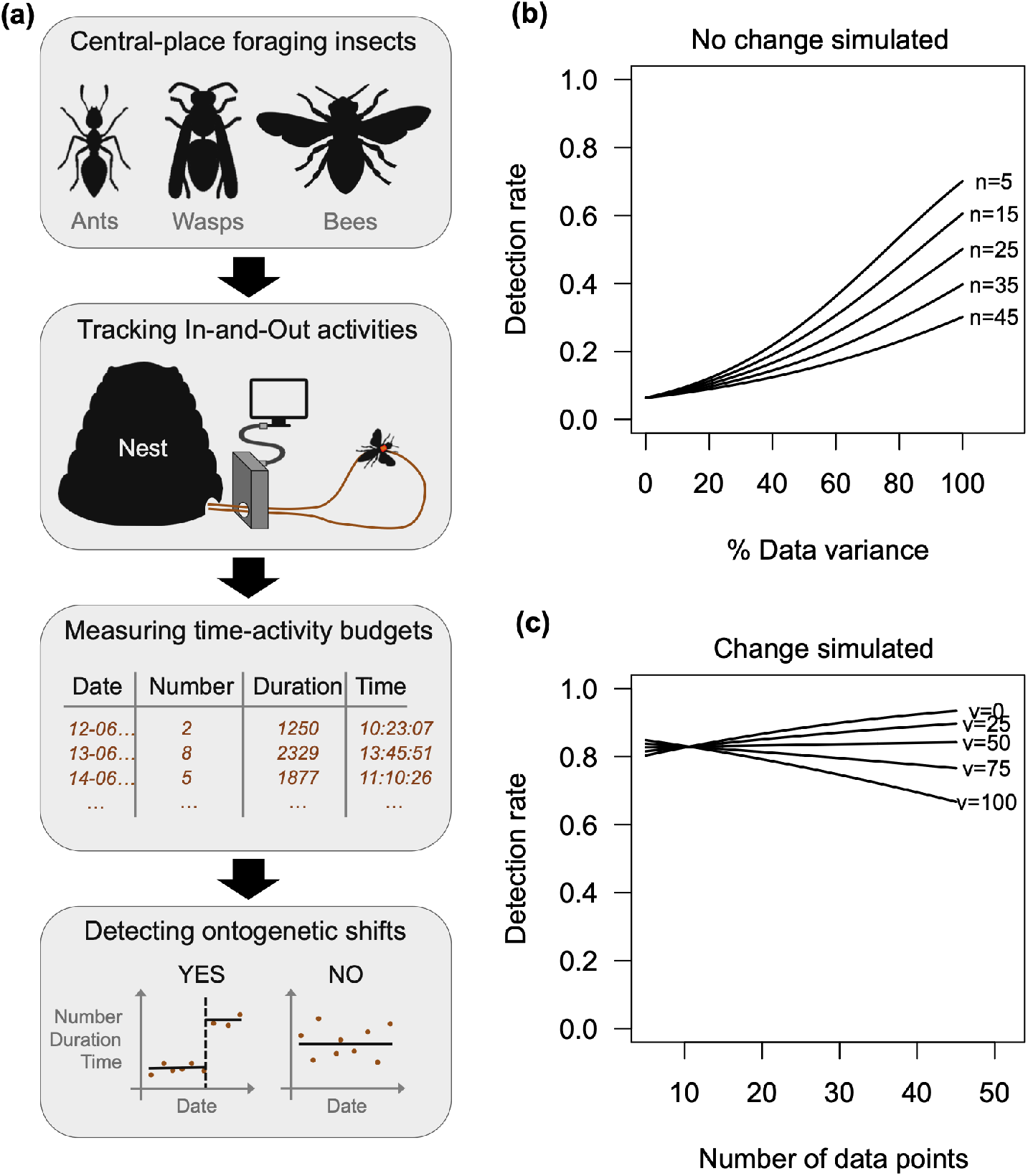
Guidelines for the use of the *aof* R-package as an automated breakpointbased procedure to detect ontogenetic shifts in central-place foraging insects. (a) The method was initially developed to detect the Age at Onset of Foraging (AOF) in honey bees, nevertheless, it could be of interest for measuring other behavioural changes in other species (e.g. ants, wasps). The in-and-out activity of central-place foraging insects is now easily recorded by miniaturized life-long individual monitoring devices, producing detailed information on time-activity budgets over the complete life of insects. The *aof* procedure can detect behavioural change along time-activity budget series. Based on simulated time-series data, we assessed (b) the sensitivity of the procedure to detect behavioural changes when there are none, and (c) the robustness of the procedure to detect existing behavioural change. For simplicity of illustration, are only shown model predictions for five categories of number of data points in b and data variance in c. These simulations will help defining the range of dataset for which the *aof* procedure is robust to detect ontogenetic shifts.

## Discussion

Changes in the temporal pattern of time-activity budgets are indicative of a transition stage in ontogenetic development (Lambin & Yoccoz, 2001; Gopukumar, 2003; Benton et al., 2008; Fay et al., 2016). Life-long individual monitoring that measures time-activity budgets provides a way to quantify a shift in the temporal pattern in time-activity budgets of insects. In this study, we develop a simple statistical procedure, derived from a breakpoint-based method, to detect such shifts in central-place foraging insects when using individual life-long monitoring (**Figure 5a**). The automated method is freely available through the *aof* R-package that includes documentation, source code and examples using both simulated and empirical dataset (see **Section 2**).

The AOF ontogenetic shift developed here is consistent with previous published estimates or observations in honey bees, e.g. orientation flights are shorter in duration than foraging flights (respectively 1 – 10 min/trip versus > 10 min/trip; Thom, Seeley, & Tautz, 2000), the number of trips per day (5.1 ± 6.1: 3.5 in Thom et al. (2000)), and the duration of foraging trips (36.0 ± 21.7 minutes compared to 38 and 50 min in Fewell and Winston (1996)). We estimated an AOF of 22.4 ± 10.6 days, consistent with other estimates, e.g. 12.8 days in Dukas (2008), 14 days in Capaldi et al. (2000), 20.6 days in Seeley (1982), and 25.6 days in Winston and Punnett (1982). Beyond ontogenetic shifts, our individual life-long monitoring also allows collecting other life-history traits, such as in-nest tasks (12.2 ± 6.9 days), which was consistent with 6 days in Capaldi et al. (2000), 8.9 days in Seeley (1982), but lower than 25.7 days in Winston and Punnett (1982). The lifespan was 31.1 ± 12.5 days in our study, also matching with previous estimates: 30 days in Schmid-Hempel and Wolf (1988), 19.9 – 25.9 days in Becerra-Guzman et al. (2005), 33.0 – 38.4 days in Amdam, Rueppell, Fondrk, Page, and Nelson (2009). Finally, our estimate of 8.7 ± 4.1 days for the duration of the foraging task was similar to other published data: 7.7 days in Visscher and Dukas (1997), and 6.8 days in Dukas (2008). Given that the foraging stage varied, we found that the AOF ontogenetic shift affected this foraging performance, with an intermediary optimal value (about 41 days). These results confirm that a precocious transition from the learning stage to the foraging stage will result in a weakened foraging performance of the honey bees (Visscher & Dukas, 1997; Becerra-Guzman et al., 2005; Rueppell et al., 2007; Perry et al., 2015). Although values of ontogenetic shifts seem realistic in comparison with previous work, they are limited to the context of the study (i.e. few colonies and a low number of different environments). The main outcome of the study is the approach exploring from data whether ontogenetic shifts in activity patterns can be detected. This analytic approach will allow further studies assessing ontogenetic shifts from data with large number of colonies and diverse environments.

The robustness assessment confirmed the ability of the breakpoint-based method to estimate the AOF ontogenetic shift in honey bees. While a previous experiment confirmed the high detection rate of our individual life-long monitoring device on honey bees (**Section S4**), tracking insects with data loggers is still challenging (Kissling et al., 2014) with a common occurrence of missed detections (Chen et al., 2012; Alaux, Crauser, Pioz, Saulnier, & Le Conte, 2014; Ngo et al., 2019 for image-based optical counter applications, and for RFID see Streit et al., 2003; Decourtye et al., 2011; Henry et al., 2012; Tenczar et al., 2014; He et al., 2015; Perry et al., 2015), that can bias the record of time-activity budgets. We recommend using the simple hit-based AOF assessment method whenever the detection rate is suspected to be low (e.g. < 95 %) and when such hit data are available (e.g. not systematically recorded with image-based optical counters; Prado et al., 2019). This simple hit-based method offers a good compromise between data management complexity and uncertainty. It relaxes the assumption of unaltered trip sequences, while enabling AOF assessment in a greater number of individuals than the simple threshold-based methods. Whenever the detection approaches 100 %, the trip-based AOF assessment would be recommended using the *Trip duration* breakpoint-based method (or the combination of *Trip duration* and *Trip time* methods given the marginally significant effect on the latter). Although the age at onset of foraging appeared to be highly plastic among bees, this simple statistical procedure was able to detect it, conversely to arbitrary thresholds that ignore individual variability.

Western honey bee colonies are currently suffering collapse disorder (Potts et al., 2010; Goulson, Nicholls, Botías, & Rotheray, 2015), a phenomenon that has motivated empirical studies using individual life-long monitoring to assess the impact of environmental stressors on the life-history of forager bees and colony fitness. For instance, several studies have suggested that the life-history of honey bees could be disturbed by pesticide exposure (e.g. Henry et al., 2012), invasive predators (e.g. Arca et al., 2014; Requier et al., 2019) and scarcity of floral resources (e.g. Requier, Odoux, Henry & Bretagnolle, 2017). Other studies further proposed that anthropogenic disturbances and their impacts on life-history traits could directly affect the survival probability of honey bee colonies (Henry et al., 2012; Becher et al., 2014; Perry et al., 2015; Requier et al., 2017). We believe that the recording of the AOF ontogenetic shift could represent an early-warning indicator of the risk of colony collapse, as previously suggested by Perry et al. (2015) for which similar effects of AOF on foraging performance have been shown. Recording the life-history and ontogenetic shifts in honey bees at the individual level and in field-realistic conditions should therefore represent a priority in the current context of global decline of bee populations (Goulson et al., 2015; Perry et al., 2015). It can then be used as an empirical baseline for the modelling of the honey bee colony (i.e. multi-agent system and mechanistic model, see Becher et al., 2014; Henry et al., 2017) with more accurate estimations of life-history parameters including and accounting for behavioural plasticity. Consequently, our method could help the epidemiological surveys to measure the impact of current environmental stressors on honey bee ecology in farmland habitats, such as pesticide intoxication, expansion of invasive predators and scarcity of floral resources (Goulson et al., 2015).

The *aof* automated method using individual life-long monitoring is generic enough to be applicable to any tracking devices that record time-activity budgets in free-ranging insects (i.e. harmonic radar, image-based optical counter, and RFID). While developed for the study on honey bees, the present method could propose new ecological applications of individual life-long monitoring devices to detect ontogenetic shifts in the behaviour of other central-place foraging insects, such as bumble bees (Gill & Raine, 2014; Crall et al., 2018; Minahan & Brunet, 2018), wasps, hornets (Poidatz, Monceau, Bonnard, & Thiéry, 2018) and ants (Pinter-Wollman et al., 2012). The sensitivity analysis showed a higher performance of the *aof* method to detect behavioural changes when the number of data points in time series increased, and an additional performance gain whenever the data variance decreased. These data criteria need to be considered as guidelines for detection of ontogenetic shifts in central-place foraging insects.

## Authors’ contributions

F.Req. and V.B. conceived the ideas and designed methodology; F.Req. collected and analysed the data; F.Req. led the writing of the manuscript and V.B., M.H., A.D., F.B., P.A. and F.Reb. contributed critically to the drafts. All authors gave final approval for publication.

## Acknowledgments

This work was supported by grants from the French Ministry of Agriculture (CASDAR, POLINOV project n°9535), the Poitou-Charentes Region and the European Community program (797/2004) for French beekeeping coordinated by the French Ministry of Agriculture (TECHBEE and RISQAPI projects). Special thanks to Jean-François Odoux and Clovis Toullet for logistical help in fieldwork. We are grateful to Cynthia McDonnell for critical reading of English used in the text, as well as the associate editor and three anonymous reviewers for constructive comments on the manuscript.

## Data accessibility

Documentation and source code to detect ontogenetic shifts in central-place foraging insects are freely available on CRAN (https://cran.r-project.org/package=aof) and GitHub (https://github.com/frareb/aof). The *aof* R-package can be installed in R with the following command: install.packages(“aof”). Moreover, data on life-history traits of honey bees are publically available through the figshare repository (doi: xx.xxxx/xx.fidshare.xxxxxxxx.xx [to be complete before final acceptance]).

