## Supporting Information for "Measuring ontogenetic shifts in central-place foraging insects: a case study with honey bees"

Requier et al.

**SUPPORTING INFORMATION**

| **Content** |  | **pj** |
| --- | --- | --- |
| Figure S1 | Location of the study sites and land-use composition over the surrounding landscapes. | 2 |
| Figure S2 | Conceptual diagram of life-history of honey bee workers. | 3 |
| Section S1 | Measure of time-activity budgets of honey bees with common 10-frame hives. | 4 |
| Section S2 | Using the *aof* R-package to measure ontogenetic shifts in central-place foraging insects | 7 |
| Section S3 | Pairwise comparisons between candidate time-activity budget variables. | 25 |
| Section S4 | Estimation of the missing detections. | 28 |
| Table S1 | Sample sizes. | 29 |
| Table S2 | Summary of the *t*-test comparisons of foraging time-activity budgets between confirmed foragers and presumed foragers. | 30 |
| Table S3 | Summary of the binomial GLMs performed for the sensitivity analysis. | 31 |
| **References** |  | **32** |

**Figure S1.** Location of the study sites and land-use composition over the surrounding landscapes.

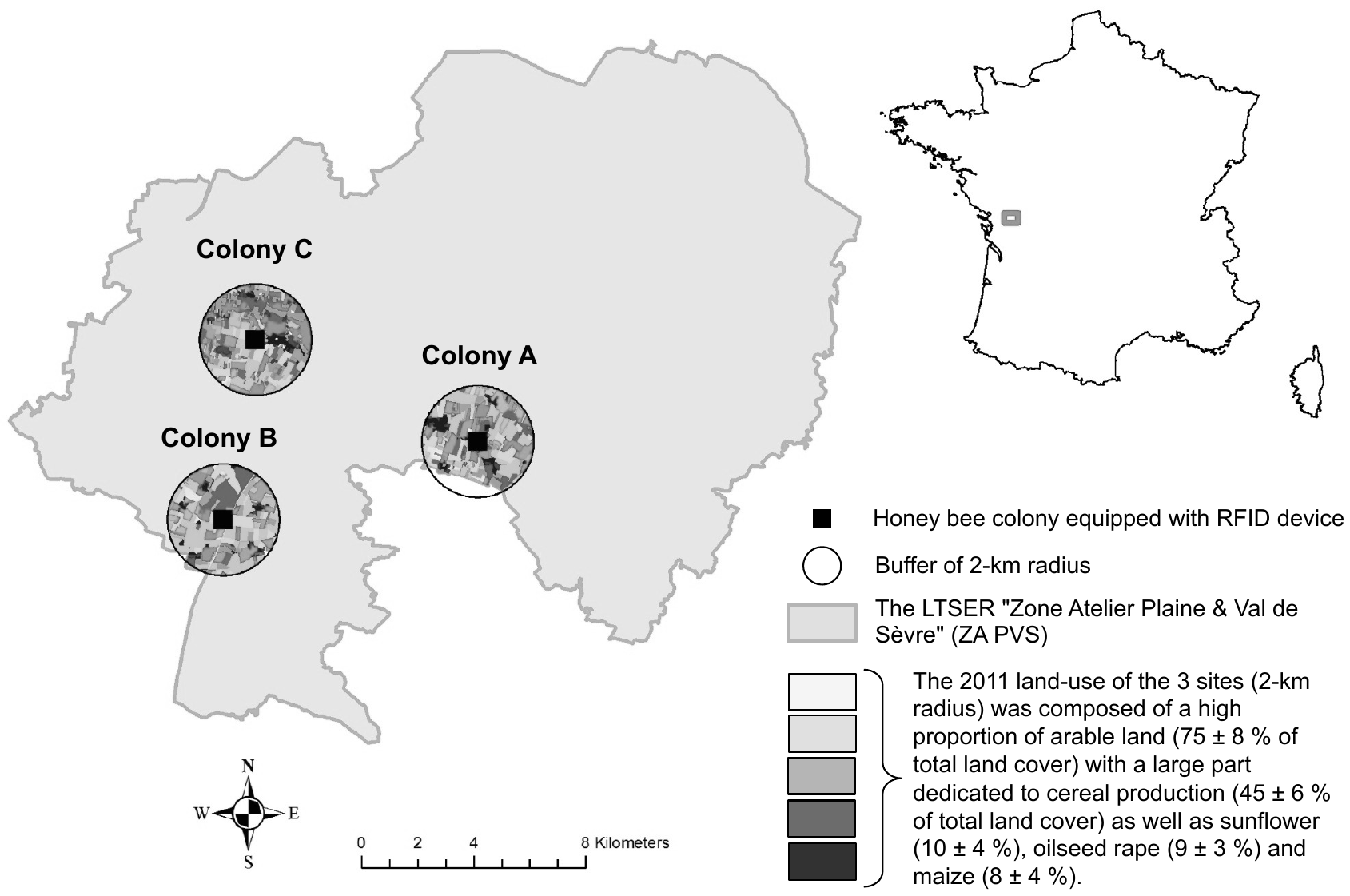

**Figure S2. Conceptual diagram of the life-history of honey bee workers.** We presented the estimated value of each age of task transition, i.e. B = Birth; AFE = Age of First Exit; AOF = Age at the Onset of Foraging; LSP = Lifespan, and their respective task durations, i.e. in-nest task stage before AFE; learning flight stage before AOF; foraging stage until LSP (mean ± SE). All bees did not perform all tasks (i.e., in-nest tasks + learning + foraging): 31.1 % of the monitored bees did not perform outside tasks (i.e. only in-nest tasks, see dotted arrow). Among bees performing outside tasks, in 74.3 % of cases, AOF was successfully detected (see thick black arrow, compared to 25.7 % of unsuccessful detections in grey arrow).

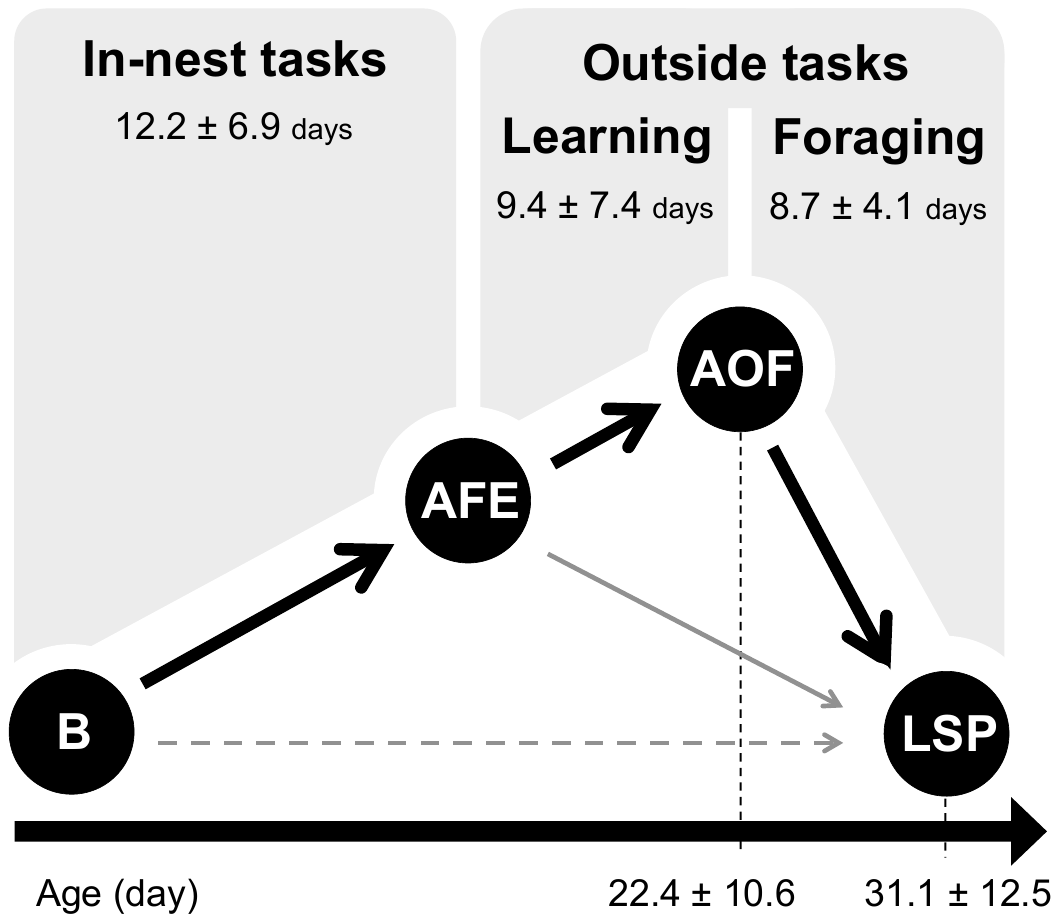

**Figure S3.** **Sensitivity analysis of the *aof* procedure based on simulated time series data.** We simulated time-series data along a factorial design for which we sequentially increased the number of data points (*n*) and the data variance (*v*). We duplicated these simulations in two sets of scenarios, (**a**) a “no change simulated” scenario to assess the sensitivity of the procedure to detect behavioural changes when there are none, and (**b**) a “change simulated” scenario to assess the robustness of the procedure to detect existing behavioural change. Grey polygons in **b** show the true value of detection, corresponding to the range of two closer data points around day 25 (the simulated breakpoint).

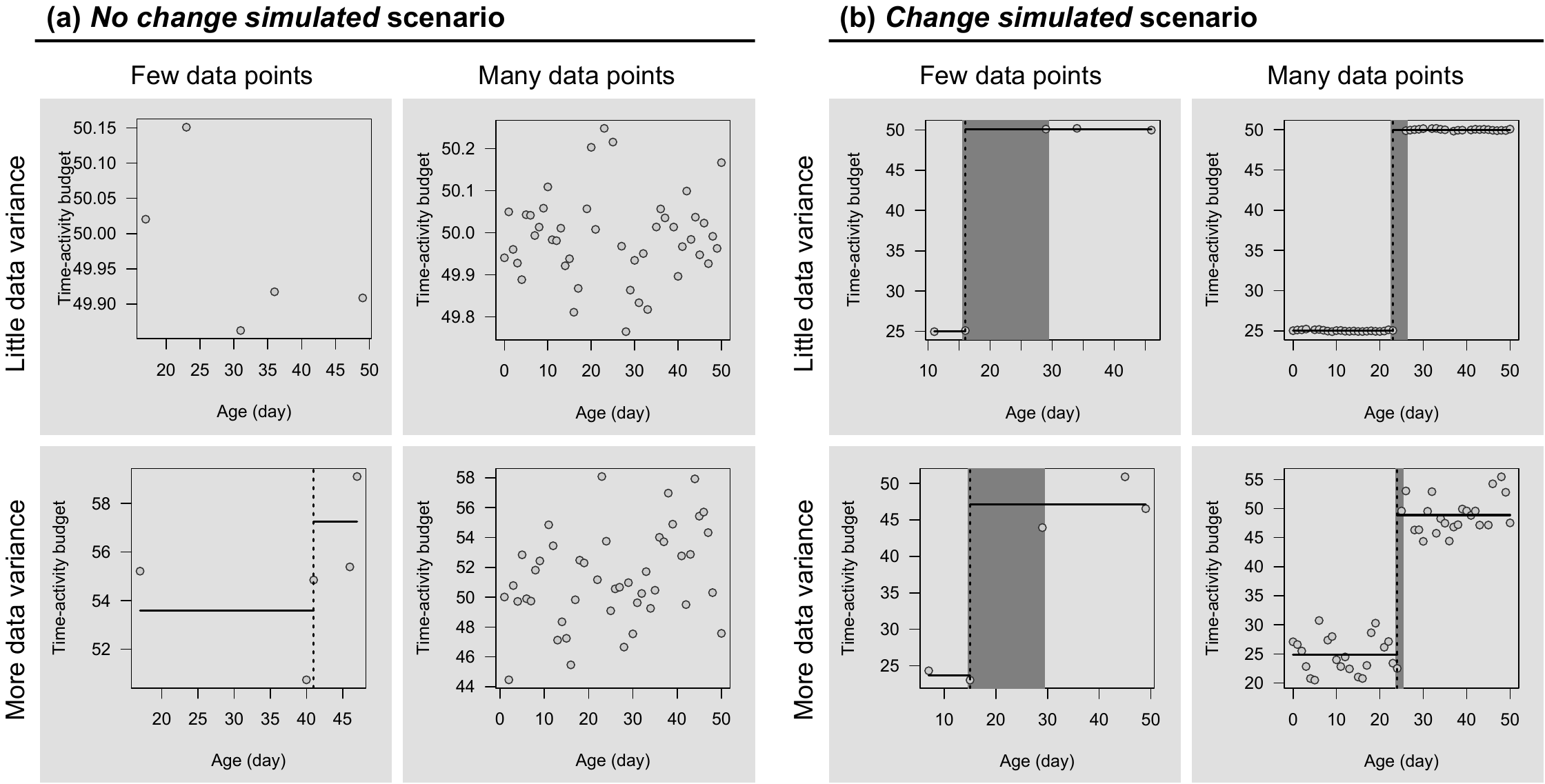

**Section S1.** Measure of time-activity budgets of honey bees in common 10-frame hives.

We used a similar approach to “Track a forager” (Van Geystelen et al., 2015) to decode the row RFID data to time budget activities. However, our specific experimental setup (i.e. establishment of two rows of five RFID readers over the entire width of the beehive entrance) discards the default setting of “Track a forager” for which we added some adaptations. We present here a meticulous step-by-step process of RFID data management to translate simple *hit* series into time budget activities (**Figure A**).

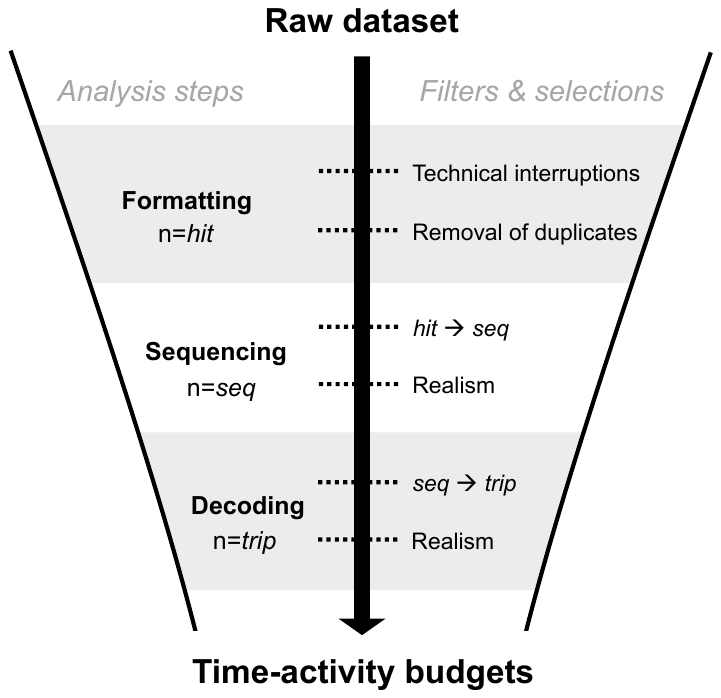

**Figure A:** Illustration of the step-by-step process to infer time-activity budgets from raw RFID dataset.

The first step consisted of data formatting, i.e. preparing the raw data files into a format readily usable for time-budget analyses. This includes the identification of *NA* (not available) time series from the monitoring database, induced by either voluntary or technical disruptions due to maintenance or power disruption. Those blackout periods (about three days out of eight months in the present study) need be explicitly distinguished from true zeroes due to the absence of flight activity. In addition, a single bee passage under a reader normally produces several successive hits due to the high recording frequency. One therefore needs to merge multiple successive hits into a single event record. We empirically established that a 1-s maximal duration interval between successive hits was a well-suited criterion for hit merging.

The second step, sequencing, consisted of translating the collected hits into sequences *seq* of I and O in order to deduce the directionality of events between the inner and outer reader rows. The following example shows a raw sequence of successive hits recorded for the tag ID #*A00103C00020BF30*:

III**IOOI**IIII**IOOOI**IOIOOI**IOI**IIIIOIOIOOOOIIIIOOIIIIOOIIIOOIIOOOIIIIIOOIOOOIIOIIIIIOOOIIOIIOOIIIIOIOIOOIOOOOIIIIOOIIOOOIIOIIOOIIIOOIOIIII

A typical flight away from the colony should return the succession IOOI, with IO giving the outgoing event from inner to outer reader rows and OI, the incoming event from outer to inner rows. Abnormal deviations from this basic pattern include elongated OO chains, e.g. IOOOOI, meaning the tagged bee stayed a while in the intra-IO free-walking room (**Figure A**), between O and I rows and then left for a second outgoing trip before the final incoming event OI. Reciprocally, elongated II chains indicate that the tagged bee has passed back and forth between the in-hive environment and the intra-IO free-walking room. Translating elongated OO or II chains into incoming and outgoing trips first and foremost depends on whether the intra-IO free-walking room should be viewed as the inside or outside of the hive. In this study, both views may be supported. Although the intra-IO free-walking room is external to the main hive chamber, it is designed as a free-walking space communicating among the 10 readers where social interactions and food exchange through trophallaxis are commonly observed due to the transparent roof (F. Requier, A. Decourtye; *personal observations*). As a parsimonious sequencing rule, i.e. delivering the theoretically smallest number of trips, we merged elongated OO chains into a single trip (rule C in **Figure B**). Two less parsimonious rules (A and B in **Figure B**) were also computed to cover a broader range of alternatives, rule A returning the maximal theoretical number of trips, and rule B an intermediate number. In all cases, elongated II chains were interpreted as a single in-hive resting bout. Abnormal records also include odd chain lengths. In the odd O-chain IOOOI, an O-hit is theoretically missing most probably due to an undetected bee passage or an unusually slow bee passage that is inadequately processed with the 1-s criterion for hit merging. Odd chains were supplemented by an additional virtual hit, and processed as normal even chains. For the sake of simplicity in the remainder of the study, statistics on AOF computation (the age at onset of foraging) with trip duration and trip time based on sequencing methods A and B are not shown. Though less parsimonious than method C in terms of trip numbers, they returned fairly similar AOF estimates (F. Requier; *personal observations*).

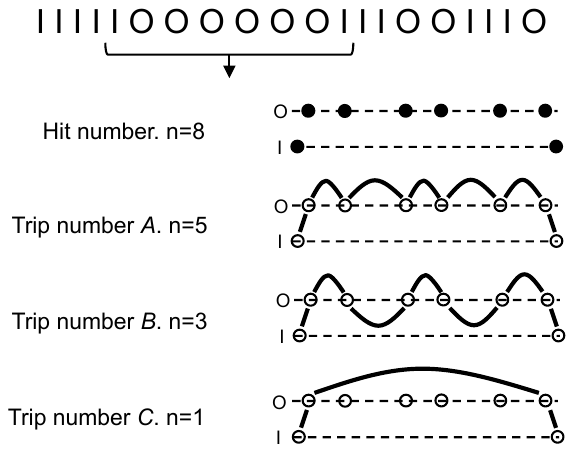

**Figure B.** Illustration of the abnormal deviations in the sequencing phase due to the specificity of our individual monitoring device adapted to a common Dadant beehive in which two rows (I and O) of five RFID readers, span the entire width of the hive entrance with a free-walking room between these two rows of readers.

As a final step of RFID data treatment, decoding, we focused only on the out activity of the bees (i.e., OO chains), reflecting the investment of bees in flight-learning (including hygienic, guarding, and orientation flights; Seeley, 1982) and foraging tasks. Unrealistic trip durations were discarded from the dataset. We set the minimal trip duration at 4-s, i.e. twice the lag time between detection and take-off, and the maximal duration as 2-h, i.e. more than 10 times the average duration of normal flights (estimated around 5 – 20 minutes according to Winston & Katz, 1982; Dukas & Visscher, 1994; Capaldi et al., 2000). Such a ceiling threshold is large enough to permit the study of adaptive flight plasticity while setting apart incoherent values due to weather disturbances or the so-called drift of bees between hives.

**Section S2.** Using the *aof* R-package to measure ontogenetic shifts in central-place foraging insects

The *aof* R-package help detecting ontogenetic shifts in univariate time-activity budget series of central-place foraging insects. The method was developed to detect the Age at Onset of Foraging (AOF) in honey bees, but can be used for the detection of other ontogenetic shifts in other central-place foraging insects. To facilitate the use of this procedure by other researchers, this **Section S2** provides documentation, source codes and examples with both simulated and empirical dataset. More documentation is also freely available on CRAN (https://cran.r-project.org/package=aof) and GitHub (https://github.com/frareb/aof). The *aof* R-package can be installed in R (R Development Core Team, 2020) with the following command: install.packages(“aof”). Otherwise, it can be also installed from GitHub with the following command : install.packages("devtools") devtools::install_github("frareb/aof")

Example

```
library("aof")
library("bcpa")
#> Loading required package: Rcpp
#> Loading required package: plyr
```

A breakpoint-based method to detect ontogenetic shifts in univariate time-activity budget series of central-place foraging insects. The method finds a single changepoint in time series where parameters change at some unknown timepoints t* is done by simply sweeping all possible breaks, and finding the most-likely changepoint according to the likelihood. The method was developed with honey bees in order to detect the Age at Onset of Foraging (AOF), but can be used for the detection of other ontogenetic shifts in other central-place foraging insects. For more details, see Requier et al. (2020) Measuring ontogenetic shifts in central-place foraging insects: a case study with honey bees. Journal of Animal Ecology.

```
# mu1 and mu2: behavioural values at stage 1 and stage 2.
# Both values mu1 and mu2 are equal (e.g. mu1=mu2=50) if no behavioural change is
# simulated, or different (e.g. mu1=25 and mu2=50) if behavioural change is
# simulated.
# rho1 and rho2 : interval frequency (default value 0.5 for both stages)
# n.obs: no. observations randomly selected in the time series, from 5 to 45
# sigma1 and sigma2: variance around the behavioural value, from 0.1 to 3
# t.full: time series from 0 to 50
# n.obs: no. observations randomly selected in the time series, from 5 to 45
# t.break: the time of the simulated behavioural change (default value of 25)

getTimeBudget <- function(
  mu1 = 50,
  mu2 = 50,
  rho1 = 0.5,
  rho2 = 0.5,
  n.obs = 5,
  sigma1 = 3,
  sigma2 = 3,
  t.full = 0:50,
  t.break = 25
){
  SimTS <- function(n, mu, rho, sigma){
    X.standard <- arima.sim(n, model = list(ar = rho))
    X.standard/sd(X.standard)*sigma + mu
  }
  x.full <- c(SimTS(t.break, mu1, rho1, sigma1),
    SimTS(max(t.full) - t.break + 1, mu2, rho2, sigma2))
  keep <- sort(sample(1:length(x.full), n.obs))
  TimeBudget <- data.frame(
    name = "A",
    Age = t.full[keep],
    x = x.full[keep])
  return(TimeBudget)
}
getAofPlot <- function(
  TimeBudget = TimeBudget,
  AOF = AOF,
  t.break = 25,
  poly = FALSE,
  ylabX = "x"
){
  par(mar = c(4, 4, 1, 1), mfrow = c(1, 1))
  plot(
    x = TimeBudget$Age,
    y = TimeBudget$x,
    las = 1,
    xlab = "Age (day)", ylab = ylabX,
    pch = 21,
    col = "gray20", bg = "gray80")
  goodbreak1 = max(TimeBudget$Age[TimeBudget$Age < t.break])
  goodbreak2 = min(TimeBudget$Age[TimeBudget$Age >= t.break])
  if(poly == TRUE){
    polygon(
      c(
        goodbreak1 - 0.5,
        goodbreak2 + 0.5,
        goodbreak2 + 0.5,
        goodbreak1 - 0.5),
      c(0, 0, 100, 100),
      col = "gray50", border = NA)
  }
  abline(v = AOF$AOF, col = "black", lwd = 2, lty = 3)
  lines(
    x = c(min(TimeBudget$Age), AOF$AOF),
    y = c(AOF$behav.stage1, AOF$behav.stage1),
    col = "black", lwd = 2)
  lines(
    x = c(AOF$AOF, max(TimeBudget$Age)),
    y = c(AOF$behav.stage2, AOF$behav.stage2),
    col = "black", lwd = 2)
  return(c(goodbreak1, goodbreak2))
}
```

### 1. Examples with no change simulated in the time series

#### 1.1. Low number of individuals (N, n.obs) and low variance (V, sigma)

```
TimeBudget <- getTimeBudget(
  n.obs = 5,
  sigma1 = 0.1,
  sigma2 = 0.1)
print(TimeBudget)
#>   name Age        x
#> 1    A  16 50.00586
#> 2    A  18 50.13609
#> 3    A  20 50.28156
#> 4    A  32 50.04075
#> 5    A  44 50.09769
AOF <- aof(
  name = TimeBudget[,1],
  Age = TimeBudget[,2],
  x = TimeBudget[,3])
#>   |                                                                              |                                                                      |   0%  |                                                                              |======================================================================| 100%
print(AOF)
#>   name parameter        AOF behav.total behav.stage1 behav.stage2
#> 1    A         x undetected    50.11239           NA           NA
getAofPlot(TimeBudget, AOF)
```

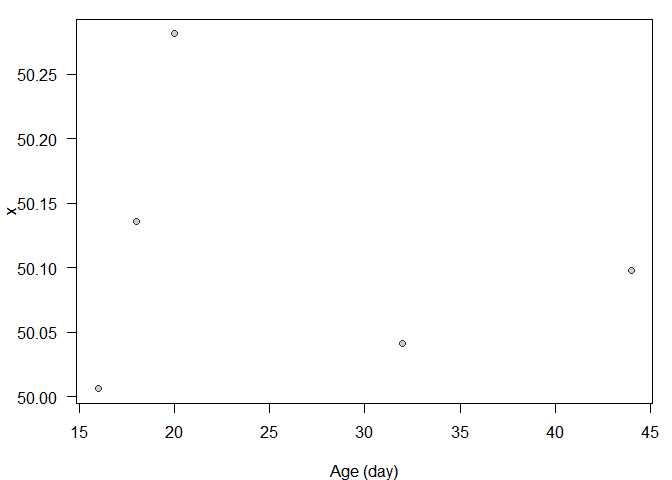

#> [1] 20 32

#### 1.2. Low number of individuals and high variance

```
TimeBudget <- getTimeBudget(
  n.obs = 5,
  sigma1 = 3,
  sigma2 = 3)
print(TimeBudget)
#>   name Age        x
#> 1    A   3 53.74759
#> 2    A   7 48.53779
#> 3    A  14 52.12544
#> 4    A  20 54.09422
#> 5    A  45 44.79826
AOF <- aof(
  name = TimeBudget[,1],
  Age = TimeBudget[,2],
  x = TimeBudget[,3])
#>   |                                                                              |                                                                      |   0%  |                                                                              |======================================================================| 100%
print(AOF)
#>   name parameter AOF behav.total behav.stage1 behav.stage2
#> 1    A         x  14    50.66066     51.47027     49.44624
getAofPlot(TimeBudget, AOF)
```

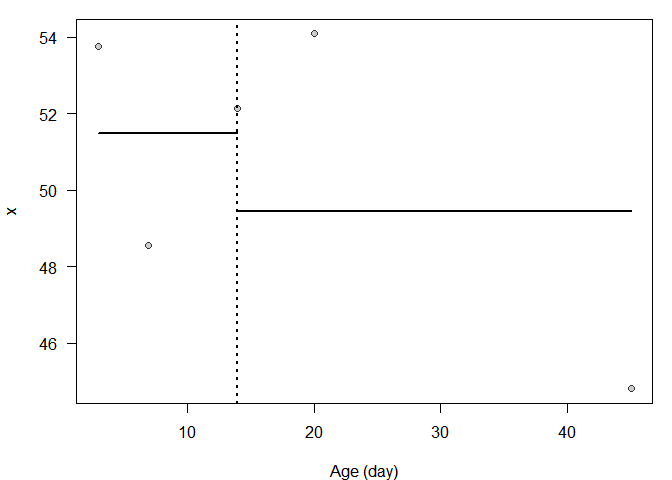

#> [1] 20 45

#### 1.3. High number of individuals and low variance

```
TimeBudget <- getTimeBudget(
  n.obs = 45,
  sigma1 = 0.1,
  sigma2 = 0.1)
print(TimeBudget)
#>    name Age        x
#> 1     A   0 49.95172
#> 2     A   1 50.06851
#> 3     A   2 50.16754
#> 4     A   3 50.11726
#> 5     A   4 50.05397
#> 6     A   5 50.16350
#> 7     A   6 50.08584
#> 8     A   7 50.06759
#> 9     A   8 49.96344
#> 10    A  11 49.89160
#> 11    A  12 49.87577
#> 12    A  13 49.97889
#> 13    A  15 49.91669
#> 14    A  16 49.91652
#> 15    A  17 49.88225
#> 16    A  18 49.82374
#> 17    A  19 49.86812
#> 18    A  20 49.99937
#> 19    A  21 49.97393
#> 20    A  23 50.00978
#> 21    A  24 50.15591
#> 22    A  25 49.95495
#> 23    A  26 50.01495
#> 24    A  27 50.02719
#> 25    A  28 50.00995
#> 26    A  29 50.13994
#> 27    A  30 50.16789
#> 28    A  31 50.01537
#> 29    A  32 49.92839
#> 30    A  33 50.01842
#> 31    A  34 50.01806
#> 32    A  35 49.94948
#> 33    A  36 50.12321
#> 34    A  37 49.94281
#> 35    A  38 50.25392
#> 36    A  39 50.06124
#> 37    A  40 49.84819
#> 38    A  41 50.03910
#> 39    A  43 50.10524
#> 40    A  44 50.16737
#> 41    A  46 49.88773
#> 42    A  47 50.06026
#> 43    A  48 50.16283
#> 44    A  49 50.02310
#> 45    A  50 50.06355
AOF <- aof(
  name = TimeBudget[,1],
  Age = TimeBudget[,2],
  x = TimeBudget[,3])
#>   |                                                                              |                                                                      |   0%  |                                                                              |======================================================================| 100%
print(AOF)
#>   name parameter        AOF behav.total behav.stage1 behav.stage2
#> 1    A         x undetected    50.02034           NA           NA
getAofPlot(TimeBudget, AOF)
```

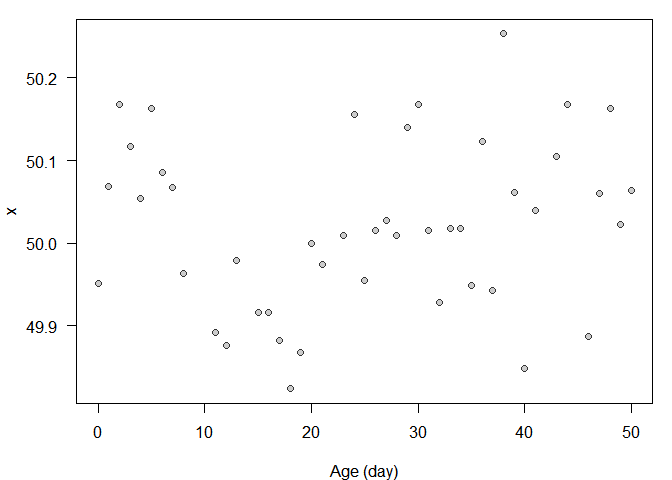

#> [1] 24 25

#### 1.4. High number of individuals and high variance

```
TimeBudget <- getTimeBudget(
  n.obs = 45,
  sigma1 = 3,
  sigma2 = 3)
print(TimeBudget)
#>    name Age        x
#> 1     A   0 52.15012
#> 2     A   1 51.91318
#> 3     A   2 50.55088
#> 4     A   3 51.92957
#> 5     A   4 47.51636
#> 6     A   5 51.17151
#> 7     A   8 46.18511
#> 8     A   9 45.59024
#> 9     A  10 53.64229
#> 10    A  12 48.12553
#> 11    A  13 42.92231
#> 12    A  14 49.26348
#> 13    A  15 52.68304
#> 14    A  16 53.83803
#> 15    A  17 53.61325
#> 16    A  18 49.82302
#> 17    A  19 51.45674
#> 18    A  20 46.24448
#> 19    A  21 49.60735
#> 20    A  22 49.41577
#> 21    A  23 50.29752
#> 22    A  24 45.05826
#> 23    A  25 50.43574
#> 24    A  26 46.45717
#> 25    A  28 52.22451
#> 26    A  29 46.71478
#> 27    A  30 48.80910
#> 28    A  31 48.22433
#> 29    A  32 56.35952
#> 30    A  33 55.00854
#> 31    A  34 53.17389
#> 32    A  35 50.06755
#> 33    A  36 51.46976
#> 34    A  37 52.58503
#> 35    A  38 54.91278
#> 36    A  39 53.33952
#> 37    A  41 51.46951
#> 38    A  42 49.21901
#> 39    A  44 49.62075
#> 40    A  45 51.20760
#> 41    A  46 47.23645
#> 42    A  47 50.16948
#> 43    A  48 44.35206
#> 44    A  49 46.44736
#> 45    A  50 47.71735
AOF <- aof(
  name = TimeBudget[,1],
  Age = TimeBudget[,2],
  x = TimeBudget[,3])
#>   |                                                                              |                                                                      |   0%  |                                                                              |======================================================================| 100%
print(AOF)
#>   name parameter        AOF behav.total behav.stage1 behav.stage2
#> 1    A         x undetected    50.00488           NA           NA
getAofPlot(TimeBudget, AOF)
```

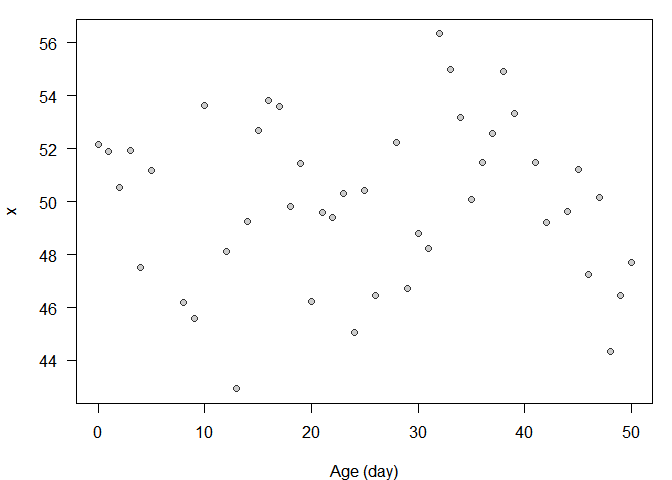

#> [1] 24 25

### 2. Examples with change simulated in the time series

#### 2.1. Low number of individuals and low variance

```
TimeBudget <- getTimeBudget(
  mu1 = 25,
  n.obs = 5,
  sigma1 = 0.1,
  sigma2 = 0.1)
print(TimeBudget)
#>   name Age        x
#> 1    A  22 25.00006
#> 2    A  23 25.00208
#> 3    A  27 49.95858
#> 4    A  37 50.01582
#> 5    A  46 49.93526
AOF <- aof(
  name = TimeBudget[,1],
  Age = TimeBudget[,2],
  x = TimeBudget[,3])
#>   |                                                                              |                                                                      |   0%  |                                                                              |======================================================================| 100%
print(AOF)
#>   name parameter AOF behav.total behav.stage1 behav.stage2
#> 1    A         x  23    39.98236     25.00107     49.96988
getAofPlot(TimeBudget, AOF)
```

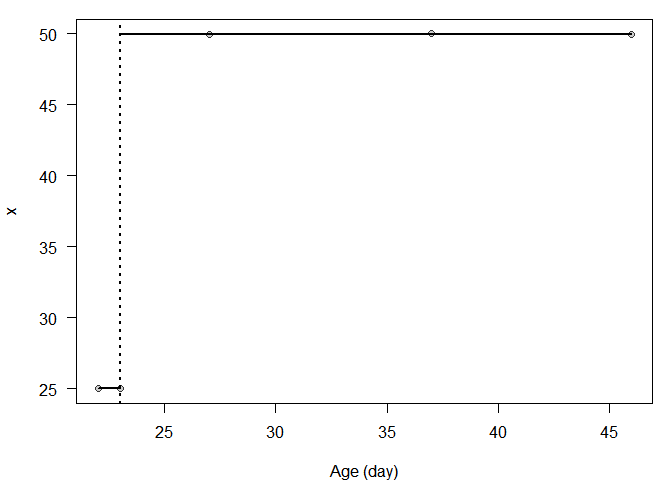

#> [1] 23 27

#### 2.2. Low number of individuals and high variance

```
TimeBudget <- getTimeBudget(
  mu1 = 25,
  n.obs = 5,
  sigma1 = 3,
  sigma2 = 3)
print(TimeBudget)
#>   name Age        x
#> 1    A   5 24.35603
#> 2    A  22 28.34613
#> 3    A  27 42.95427
#> 4    A  29 48.86142
#> 5    A  38 50.40718
AOF <- aof(
  name = TimeBudget[,1],
  Age = TimeBudget[,2],
  x = TimeBudget[,3])
#>   |                                                                              |                                                                      |   0%  |                                                                              |======================================================================| 100%
print(AOF)
#>   name parameter AOF behav.total behav.stage1 behav.stage2
#> 1    A         x  22    38.98501     26.35108     47.40762
getAofPlot(TimeBudget, AOF)
```

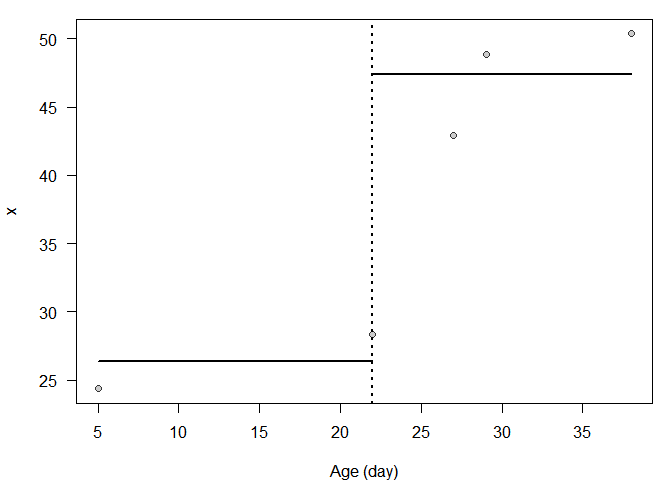

#> [1] 22 27

#### 2.3. High number of individuals and low variance

```
TimeBudget <- getTimeBudget(
  mu1 = 25,
  n.obs = 45,
  sigma1 = 0.1,
  sigma2 = 0.1)
print(TimeBudget)
#>    name Age        x
#> 1     A   0 24.95342
#> 2     A   1 24.79843
#> 3     A   3 24.79287
#> 4     A   5 24.99629
#> 5     A   6 25.11400
#> 6     A   7 25.08924
#> 7     A   9 25.05745
#> 8     A  10 25.09193
#> 9     A  12 25.04732
#> 10    A  13 24.91764
#> 11    A  15 24.85669
#> 12    A  16 24.90408
#> 13    A  17 24.83885
#> 14    A  18 24.84675
#> 15    A  19 24.90080
#> 16    A  20 25.01541
#> 17    A  21 24.91389
#> 18    A  22 24.99316
#> 19    A  23 24.94847
#> 20    A  24 24.86305
#> 21    A  25 49.96632
#> 22    A  26 49.90877
#> 23    A  27 49.85963
#> 24    A  28 50.02871
#> 25    A  29 49.87066
#> 26    A  30 50.01809
#> 27    A  31 50.06752
#> 28    A  32 50.06952
#> 29    A  33 50.06428
#> 30    A  34 49.94689
#> 31    A  35 50.00534
#> 32    A  36 49.89206
#> 33    A  37 49.90005
#> 34    A  38 49.87778
#> 35    A  39 49.92120
#> 36    A  40 49.96800
#> 37    A  41 50.02669
#> 38    A  42 49.90425
#> 39    A  43 49.84559
#> 40    A  44 50.10250
#> 41    A  45 50.16960
#> 42    A  46 50.07850
#> 43    A  48 50.11246
#> 44    A  49 50.08179
#> 45    A  50 50.16963
AOF <- aof(
  name = TimeBudget[,1],
  Age = TimeBudget[,2],
  x = TimeBudget[,3])
#>   |                                                                              |                                                                      |   0%  |                                                                              |======================================================================| 100%
print(AOF)
#>   name parameter AOF behav.total behav.stage1 behav.stage2
#> 1    A         x  26    38.86212     27.21886     49.99916
getAofPlot(TimeBudget, AOF)
```

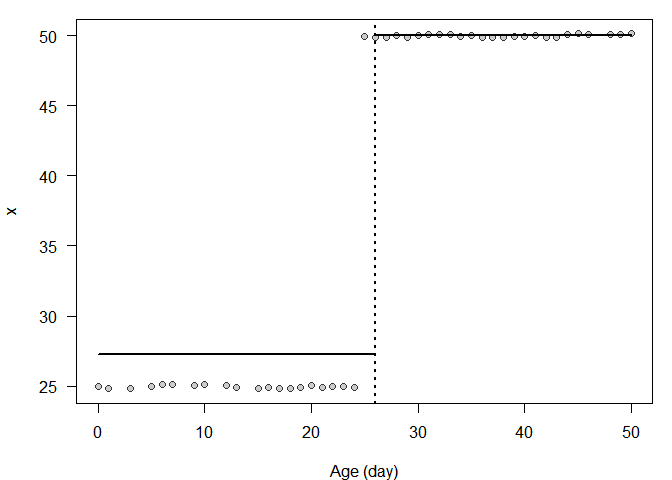

#> [1] 24 25

#### 2.4. High number of individuals and high variance

```
TimeBudget <- getTimeBudget(
  mu1 = 25,
  n.obs = 45,
  sigma1 = 3,
  sigma2 = 3)
print(TimeBudget)
#>    name Age        x
#> 1     A   0 21.40092
#> 2     A   1 21.16138
#> 3     A   3 23.57402
#> 4     A   4 26.23103
#> 5     A   6 27.47797
#> 6     A   7 31.78789
#> 7     A   8 21.56979
#> 8     A   9 29.80355
#> 9     A  10 23.38443
#> 10    A  11 28.44681
#> 11    A  12 27.52959
#> 12    A  13 26.81896
#> 13    A  14 23.28048
#> 14    A  16 23.99062
#> 15    A  17 26.18801
#> 16    A  18 23.16303
#> 17    A  19 20.21720
#> 18    A  20 21.71164
#> 19    A  21 26.60556
#> 20    A  22 24.99505
#> 21    A  23 22.93847
#> 22    A  24 24.19659
#> 23    A  25 50.01116
#> 24    A  26 51.24340
#> 25    A  27 58.59474
#> 26    A  28 55.10758
#> 27    A  29 48.67596
#> 28    A  30 45.82812
#> 29    A  31 45.44915
#> 30    A  32 51.69036
#> 31    A  33 53.14876
#> 32    A  34 51.33613
#> 33    A  37 52.81819
#> 34    A  38 52.95899
#> 35    A  39 52.93804
#> 36    A  40 47.02771
#> 37    A  42 51.80734
#> 38    A  43 50.34804
#> 39    A  44 50.60444
#> 40    A  45 49.78608
#> 41    A  46 50.44388
#> 42    A  47 46.89187
#> 43    A  48 46.72605
#> 44    A  49 46.57734
#> 45    A  50 49.35466
AOF <- aof(
  name = TimeBudget[,1],
  Age = TimeBudget[,2],
  x = TimeBudget[,3])
#>   |                                                                              |                                                                      |   0%  |                                                                              |======================================================================| 100%
print(AOF)
#>   name parameter AOF behav.total behav.stage1 behav.stage2
#> 1    A         x  24    37.90758     24.83968      50.4073
getAofPlot(TimeBudget, AOF)
```

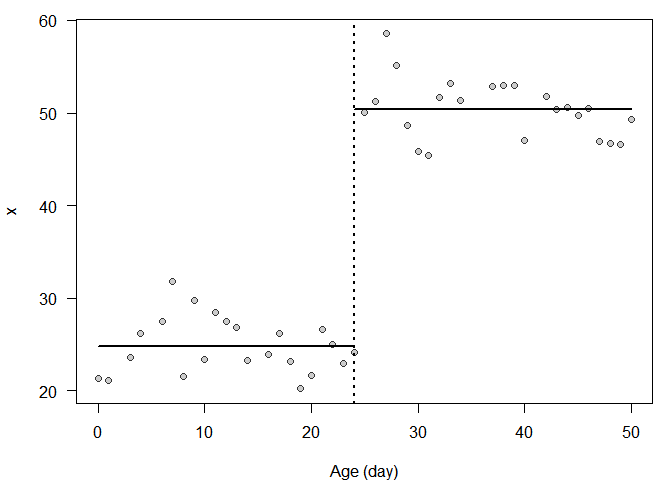

#> [1] 24 25

### 3. Real dataset

This is a subset of 5 bees randomly selected in the experimental design of Requier et al. (J. Animal Ecology).

```
TimeBudget <- dataExample
head(TimeBudget, n = 25)
#>                name Age Number  Duration    Time
#> 1  A00103C00020FD24  15      2   63.5000 55166.0
#> 2  A00103C00020FD24  20      1  176.0000 71609.0
#> 3  A00103C00020FD24  22      4 1708.5000 41340.0
#> 4  A00103C00020FD24  23      5 2762.2000 42941.0
#> 5  A00103C00020FD24  24      1  562.0000 67521.0
#> 6  A00103C00020FD24  26      2  695.0000 40153.0
#> 7  A00103C00020FD24  27      1  700.0000 48959.0
#> 8  A00103C000402590  14      1   22.0000 64725.0
#> 9  A00103C000402590  16      1  227.0000 45238.0
#> 10 A00103C000402590  23      7  205.5714 62908.0
#> 11 A00103C000402590  24     14  360.1429 54300.0
#> 12 A00103C000402590  25      3  357.3333 64024.0
#> 13 A00103C000402590  26     10  680.4000 56767.5
#> 14 A00103C000402590  27      7  781.1429 56320.0
#> 15 A00103C000402590  28      3 2786.3333 51801.0
#> 16 A00103C000402590  29      1    8.0000 42236.0
#> 17 A00103C000406D60  13      1   14.0000 56166.0
#> 18 A00103C000406D60  14      2   89.0000 60514.5
#> 19 A00103C000406D60  15      4  150.5000 46005.5
#> 20 A00103C000406D60  16      1    4.0000 68591.0
#> 21 A00103C000406D60  17      2  330.5000 58752.5
#> 22 A00103C000406D60  18      3  781.0000 54998.0
#> 23 A00103C000406D60  20      4 1981.0000 57991.0
#> 24 A00103C000406D60  22      2  735.0000 52296.5
#> 25 A00103C000406D60  23      7 1908.2857 44964.0
# working on Number of trips
varX <- "Number"
AOF_number <- aof(
  name = TimeBudget[,1],
  Age = TimeBudget[,2],
  x = TimeBudget[varX])
#>   |                                                                              |                                                                      |   0%  |                                                                              |==============                                                        |  20%  |                                                                              |============================                                          |  40%  |                                                                              |==========================================                            |  60%  |                                                                              |========================================================              |  80%  |                                                                              |======================================================================| 100%
print(AOF_number)
#>               name parameter        AOF behav.flightspan behav.learning
#> 1 A00103C00020FD24    Number undetected         2.285714             NA
#> 2 A00103C000402590    Number         23         5.222222       3.000000
#> 3 A00103C000406D60    Number         20         3.300000       2.428571
#> 4 A00103C00040A5D5    Number undetected         1.636364             NA
#> 5 A00103C00040A5F4    Number         23        11.125000       1.333333
#>   behav.foraging
#> 1             NA
#> 2       6.333333
#> 3       5.333333
#> 4             NA
#> 5      17.000000
# working on Duration of trips
varX <- "Duration"
AOF_duration <- aof(
  name = TimeBudget[,1],
  Age = TimeBudget[,2],
  x = TimeBudget[varX])
#>   |                                                                              |                                                                      |   0%  |                                                                              |==============                                                        |  20%  |                                                                              |============================                                          |  40%  |                                                                              |==========================================                            |  60%  |                                                                              |========================================================              |  80%  |                                                                              |======================================================================| 100%
print(AOF_duration)
#>               name parameter AOF behav.flightspan behav.learning behav.foraging
#> 1 A00103C00020FD24  Duration  20         952.4571      119.75000      1285.5400
#> 2 A00103C000402590  Duration  25         603.1026      234.40952      1063.9690
#> 3 A00103C000406D60  Duration  17         757.0571      117.60000      1396.5143
#> 4 A00103C00040A5D5  Duration  25         798.9091       81.64286      2054.1250
#> 5 A00103C00040A5F4  Duration  24         228.5120      211.82143       245.2026
# working on Time of trips
varX <- "Time"
AOF_time <- aof(
  name = TimeBudget[,1],
  Age = TimeBudget[,2],
  x = TimeBudget[varX])
#>   |                                                                              |                                                                      |   0%  |                                                                              |==============                                                        |  20%  |                                                                              |============================                                          |  40%  |                                                                              |==========================================                            |  60%  |                                                                              |========================================================              |  80%  |                                                                              |======================================================================| 100%
print(AOF_time)
#>               name parameter AOF behav.flightspan behav.learning behav.foraging
#> 1 A00103C00020FD24      Time  20         52527.00       63387.50       48182.80
#> 2 A00103C000402590      Time  23         55368.83       57623.67       54241.42
#> 3 A00103C000406D60      Time  18         54637.80       57504.58       50337.62
#> 4 A00103C00040A5D5      Time  25         56072.86       59164.14       50663.12
#> 5 A00103C00040A5F4      Time  25         59626.88       62104.80       55497.00
goodbreak <- lapply(seq_along(unique(TimeBudget[,1])), function(i){
  print(as.character(unique(TimeBudget[,1])[i]))
  par(mfrow = c(1, 3))
  varXList <- list("Number", "Duration", "Time")
  varAOFList <- list(AOF_number, AOF_duration, AOF_time)
  varYlabList <- list("Trip number (per day)", "Trip duration (seconds)", "Trip time (seconds)")
  varXRes <- sapply(1:3, function(j){
    TimeBx <- TimeBudget[
      TimeBudget[,1] == unique(TimeBudget[,1])[i],
      c("name", "Age", varXList[[j]])]
    names(TimeBx)[3] <- "x"
    getAofPlot(
      TimeBudget = TimeBx,
      AOF = varAOFList[[j]][i,],
      ylabX = varYlabList[[j]])
  })
})
#> [1] "A00103C00020FD24"
```

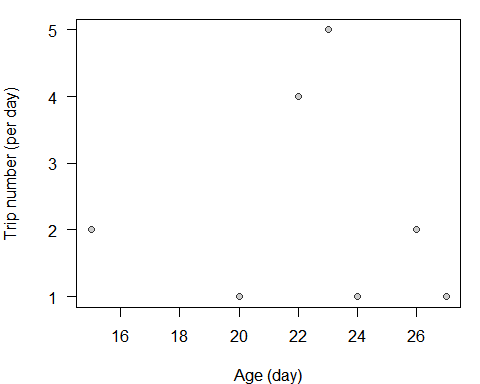

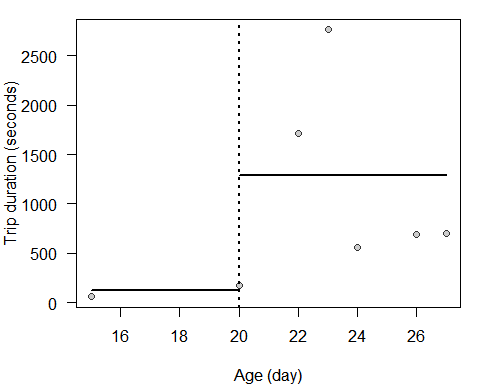

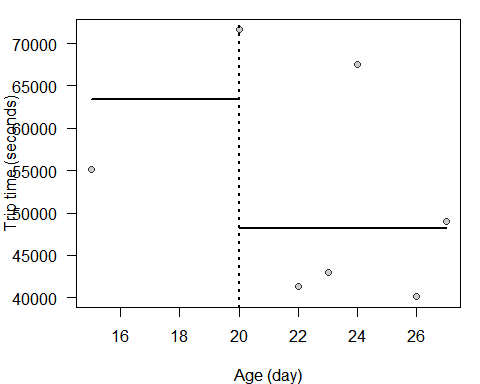

#> [1] "A00103C000402590"

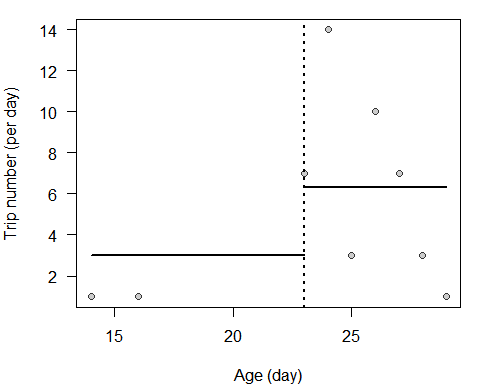

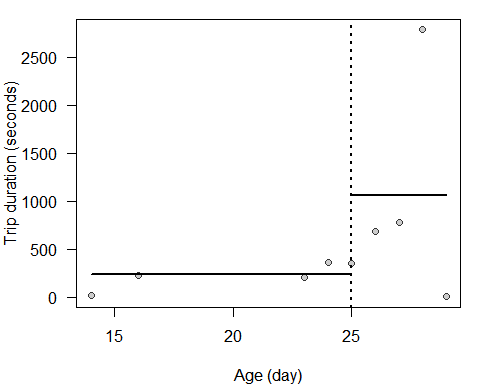

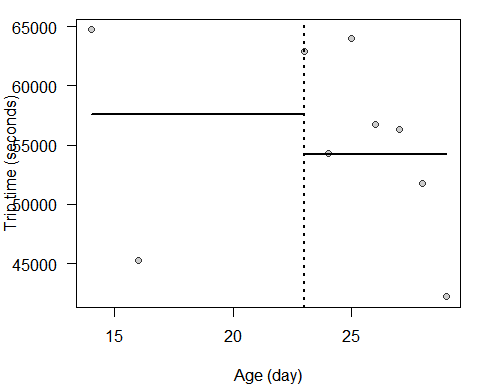

#> [1] "A00103C000406D60"

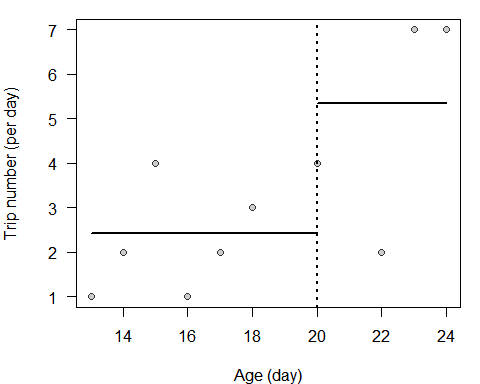

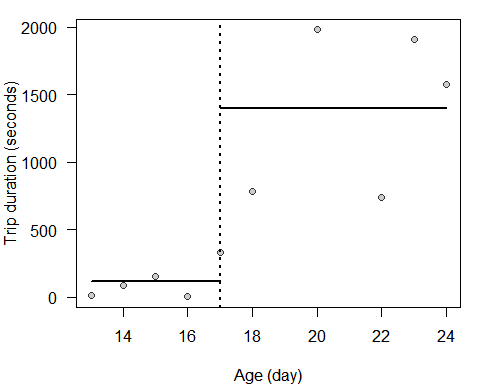

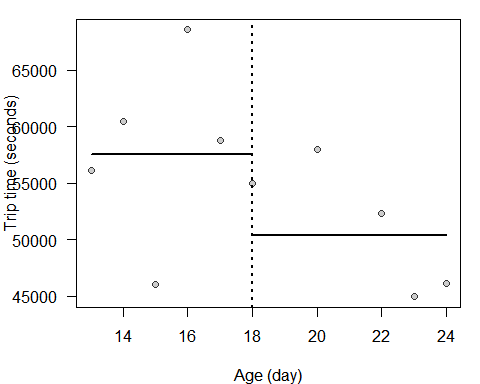

#> [1] "A00103C00040A5D5"

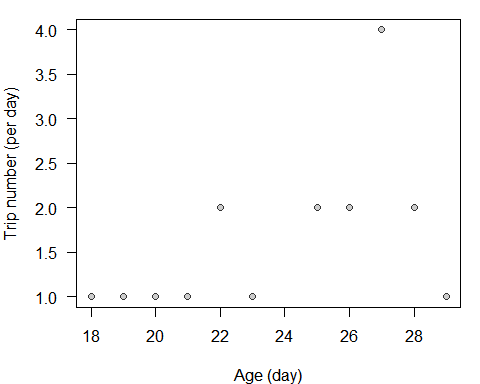

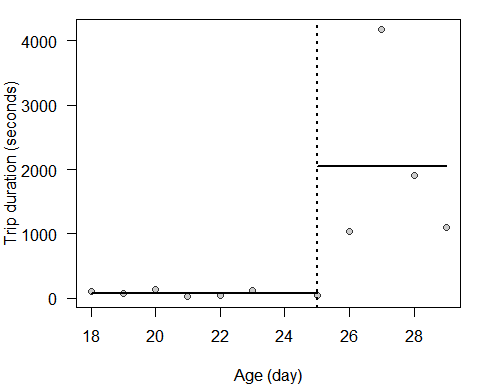

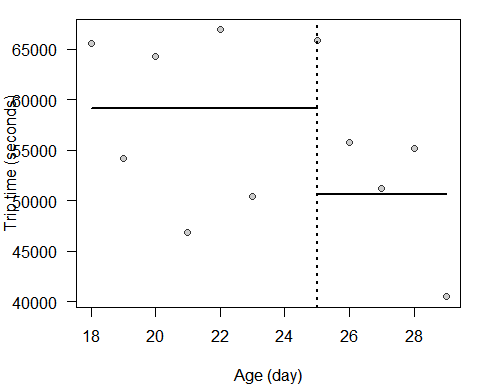

#> [1] "A00103C00040A5F4"

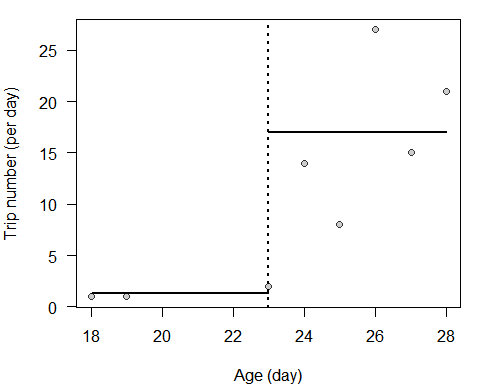

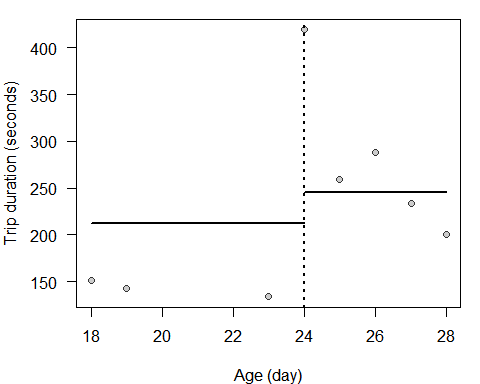

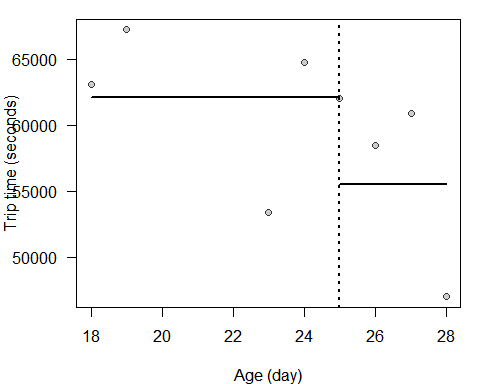

**Section S3.** Pairwise comparisons between candidate time-activity budget variables.

Pairwise comparisons were computed between each candidate time-activity budget variable: *Hit number, Hit time, Thr. trip number, Thr. trip duration, Trip number, Trip duration,* and *Trip time*. First, we estimated the pairwise differences in the value of AOF in days, at cohort level (mean ± SE). Then, we compared the number of successful AOFs detected, i.e. the size of the “detected” group. Finally, we estimated the similarity of the individuals detected, that is the proportion of shared “detected” bees between pairwise candidate time budget variables at the cohort level (crossed cohort × site grouping), and their variability (computing the coefficient of variation).

Although arbitrary, the threshold-based methods using trips detected a higher number of AOFs than breakpoint-based methods (about 778 to 983 bees *vs.* 492 to 835 bees respectively, central diagonal in **Table A**). However, the identity of those detected bees appeared to be lower between the two arbitrary threshold-based methods (64.9 %) than the similarity between the two breakpoint-based methods using hits (85.2 %) and using *Trip duration* and *Trip time* (94.8 %, **Table A**). This result suggests that threshold-based methods could be more random in the detection of AOF than breakpoint-based methods. Otherwise, breakpoint-based estimates of AOF with simple hits provided higher detection rates, from 974 to 1,114 bees, however, with detected estimates lower than other methods (**Table A**). This bias could be explained by the higher number of hits detected before the recording of the first trip (see **Figure 2b**), likely leading to early AOF estimation. All the other breakpoint-based methods (i.e. using trips) and threshold-based methods had similar estimates of AOF (**Table A**).

Surprisingly, the breakpoint-based method using *Trip number* was the less efficient technique to detect AOF (only 492 bees). In addition, the identity of those detected bees appeared to be few similar to all the other techniques; questioning on the robustness of the *Trip number* metric to detect AOF in bees. This result suggests that *Trip number* could be a bad indicator of honey bee behaviour, and should be discarded as methods to estimate AOF in bees (see also **Figure 3**).

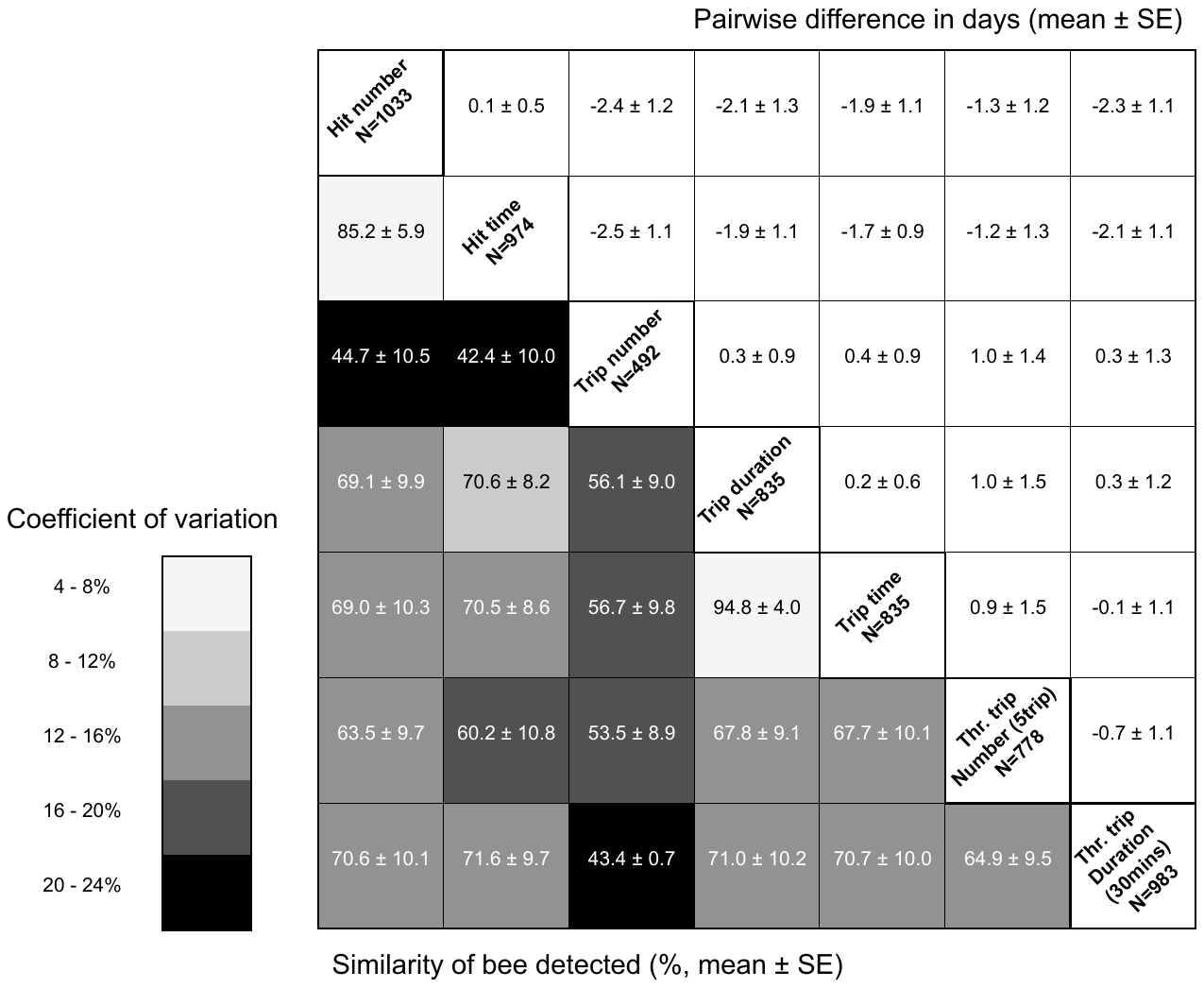

**Table A.** Pairwise comparisons between each candidate time budget variable expressed in terms of the number of successful AOFs detected (see the central diagonal, i.e. size of the “detected” group), in differences across the values of AOF in days (see top-right of the table) at cohort level (crossed cohort × site grouping, mean ± SE), and in similarity of the individuals detected (see bottom-left of the table, i.e. proportion of shared “detected” bees between pairwise variables, mean ± SE) and their variability (see the black to white gradient, computing the coefficient of variation).

**Section S4.** Estimation of the missing detections.

The sequencing algorithm is robust enough to isolate missing events but may be substantially affected by pervasive or multiple consecutive missing events. In separate laboratory trials, we assessed the detection probability of 91 tagged bees forced to walk in a tube with one-way gates and fitted with RFID readers. We obtained a satisfactory 97 % detection rate. However, experience shows that this rate may vary depending on the performance and configuration of the RFID system and on the way readers are positioned in front of the hive entrance. For the sake of comparison, Tanczar *et al.* (2014) reported a detection rate at 87.4 %.

Whenever the detection bias cannot be firmly established or is expected to be low, it might be safer to use an alternate AOF measurement approach that does not rely on the assumption of unaltered trip sequences. Herein, we suggest applying the above-described breakpoint detection procedure based on hits rather than trips. In other words, we recommend directly using the output of the formatting phase and skipping the sequencing phase (**Section S1**). Specifically, this supposes computing the daily cumulative hit numbers (*Hit number*), rather than the cumulative trip numbers, to search for breakpoints.

**Table S1.** The sample sizes for emerged honey bees and established foragers introduced into the three different colonies, with their respective dates of introductions.

| **Introduction date** | **Colony A** | | **Colony B** | **Colony C** |
| --- | --- | --- | --- | --- |
|  | Confirmed foragers | Emerged bees | Emerged bees | Emerged bees |
| 05/04/2011 | ø | 150 | 150 | ø |
| 06/05/2011 | ø | 150 | 150 | ø |
| 01/06/2011 | ø | 75 | 75 | ø |
| 21/06/2011 | 67 | ø | ø | ø |
| 30/06/2011 | ø | 150 | 150 | 150 |
| 28/07/2011 | ø | 150 | 150 | 150 |
| 25/08/2011 | ø | 150 | 150 | 150 |

**Table S2.** Summary of the *t*-test in the foraging time-activity budget allocations (i.e. number, duration and time of trips) between confirmed foragers (n = 67) and presumed foragers (n = 39 bees with AOF detected). *Thr. trip number* means the threshold-based method using 5 trips per day, and *Thr. trip duration* means the threshold-based method using flight durations longer than 30 minutes. The other time budget variables involve the breakpoint-based method on hit or trip. The Cohen’s d value compares the magnitude of difference in effect between the candidate time budget variables, ranging from 0, i.e. no difference, to 1, i.e. large difference. Bold entries indicate significant effects at a 0.05 level.

| Time-activity budget parameter | *t* | df | *P*-value | Cohen's *d*-value [95%CI] |
| --- | --- | --- | --- | --- |
| Hit time | -0.987 | 72.521 | 0.327 | -0.2 [-0.6, 0.2] |
| Trip duration | -1.451 | 52.932 | 0.153 | -0.3 [-0.71, 0.11] |
| Hit number | 1.483 | 39.548 | 0.146 | 0.3 [-0.1, 0.7] |
| **Trip time** | **-2.331** | **68.841** | **0.023** | **-0.48 [-0.89, -0.07]** |
| **Thr. trip duration** | **-2.416** | **60.509** | **0.019** | **-0.49 [-0.91, -0.08]** |
| **Trip number** | **3.299** | **26.742** | **0.003** | **0.68 [0.26, 1.09]** |
| **Thr. trip number** | **4.777** | **46.510** | **<0.001** | **0.98 [0.55, 1.41]** |

**Table S3.** Summary of the binomial GLMs performed to assess (*i*) the sensitivity of the *aof* procedure to detect behavioural change when there are none (*No change* *simulated* scenario), and (*ii*) the robustness of the procedure to detect existing behavioural change (*Change* *simulated* scenario). The “Data points” variable refers to the number of data points simulated. The s.e. refer to the standard error of model estimates. Bold entries indicate significant effects at a 0.05 level.

| ***No change* *simulated* scenario** | |  |  |
| --- | --- | --- | --- |
| Model parameter | Complete model estimate ± s.e. | *Z* | *P*-value |
| **Intercept** | **-2.689 ± 0.358** | **-7.519** | **<0.001** |
| Data points | -0.001 ± 0.013 | -0.008 | 0.994 |
| **Data variance** | **0.038 ± 0.005** | **6.980** | **<0.001** |
| **Data points × data variance** | **-0.001 ± 0.001** | **-2.120** | **0.034** |
| ***Change* *simulated* scenario** | |  |  |
| Model parameter | Complete model estimate ± s.e. | *Z* | *P*-value |
| **Intercept** | **1.250 ± 0.317** | **3.940** | **<0.001** |
| **Data points** | **0.031 ± 0.012** | **2.587** | **0.010** |
| Data variance | 0.006 ± 0.005 | 1.113 | 0.266 |
| **Data points × data variance** | **-0.001 ± 0.001** | **-2.901** | **0.004** |
